# Endocrine-enriched stem cell-derived islets improve long-term safety *in vivo*

**DOI:** 10.1101/2025.08.22.671768

**Authors:** Eunike Sawitning Ayu Setyono, Nicole Katarina Rogers, Katharina Scheibner, Silvia Schirge, Hélène Reich, Michael Sterr, Timucin Öztürk, Sören Franzenburg, Väinö Lithovius, Heiko Lickert

**Affiliations:** Institute of Diabetes and Regeneration Research, Helmholtz Diabetes Center, Helmholtz Munich, Neuherberg, Germany; German Center for Diabetes Research (DZD), Neuherberg, Germany; School of Medicine and Health, Technische Universität München, Munich, Germany; Institute of Clinical Molecular Biology, Christian-Albrechts-Universität Kiel, Kiel, Germany

**Author notes:** These authors contributed equally (Eunike Sawitning Ayu Setyono, Nicole Katarina Rogers).

## Abstract

Fully differentiated stem cell-derived islets (SC-islets) are proven to normalise blood glucose in type 1 diabetic patients. However, the presence of off-target cell types and the immature SC-islet function upon transplantation remain unresolved problems. Here, we established sorting strategies to generate SC-islets with defined glucagon-producing SC-α- and insulin-producing β-cell ratios and assessed their safety and efficacy *in vitro* and *in vivo*. Engineering SC-islets is beneficial to the insulin response *in vitro*, which does not translate to improved glycaemic regulation *in vivo*. Importantly, hormone-producing endocrine cell enrichment and thus off-target cell type depletion eliminated the risk for unwanted outgrowth *in vivo*. Single cell analysis defined off-target cells *in vitro* and *in vivo* and identified marker genes to assess SC-islet quality and define safety release criteria before graft transplantation. This study highlights the importance of determining the SC-islet composition and establishing rigorous quality controls to ensure long-term safety for β-cell replacement therapy.

## Introduction

Type 1 diabetes (T1D) is an autoimmune disease in which insulin-producing β-cells are destroyed, leading to hyperglycaemia and devastating secondary complications. Life expectancy in children diagnosed before the age of 10 is reduced by 14-18 years, with the highest risk in women^1^. While insulin replacement via injections or pump-mediated delivery remains an effective therapy for most T1D patients, a subset of them experiences frequent episodes of hypoglycaemia associated with a 3-fold increase in mortality^2^. β-cell replacement therapy by islet transplantation has the potential to cure T1D and would offer a more effective therapy, especially for patients with frequent hypoglycaemia and severe life-threatening hypoglycaemic unawareness (SHE)^3–7^, however, donor cadaveric islet availability is scarce. Thus, stem cell-derived islets (SC-islets), which can be generated in limitless quantities, have been proposed as a promising alternative cell source for islet replacement therapy for people living with diabetes^8–13^.

SC-islets are generated through a multistep differentiation process from human pluripotent stem cells (hPSC), including induced pluripotent stem cells (hiPSC) and human embryonic stem cells (hESC), by mimicking *in vivo* endoderm, pancreas and islet development^14–24^. With the successful generation of glucose-responsive SC-islets, clinical trials have been performed, from transplanting pancreatic progenitor cells^25–27^ to fully differentiated SC-islets^28,29^. Recent results in SC-islets transplantation reported improved glucose control in all twelve participants receiving full dose (800 million cells) with ten achieving insulin independence^28^.

Despite these promising functional outcomes, further optimisation is needed to ensure both efficacy and safety of SC-islets. Like primary islets, SC-islets do not exclusively consist of the insulin-producing β-cells but also contain glucagon-secreting α- and somatostatin-secreting δ-cells. α- and δ-cells have important physiological roles in counteracting hypoglycaemia^30^ and fine-tuning β-cells insulin secretion^31–36^. Their importance in the context of β-cell replacement therapy remains an open question: while maximising β-cell numbers in SC-islets could bring cost reductions by delivering the most therapeutic cells, this could come as a disadvantage for long-term impaired physiological regulation. In fact, some studies in rodents suggest that non-β-cells are necessary for long-term graft function^37^, whereas others show the opposite^38^.

SC-islets also contain cells that are not found in primary islets. These off-target cells arise as a byproduct of imperfections in the differentiation protocols and could compromise safety *in vivo* due to continued proliferation and/or differentiation^39–44^. Even SC-islets with a high proportion of endocrine cells might generate CK19^+^ cysts^43^ that could spontaneously rupture. Although the purification of SC-islets to generate β-cell-enriched clusters has shown improved insulin secretion^39,45–54^, these strategies did not address the optimal composition of SC-islets upon enrichment and the long-term effect of non-endocrine cells on functionality and safety.

Additionally, transplanting SC-islets still requires the use of immunosuppressants. In the recent phase 1-2 clinical trial, one participant had an immunosuppressant-related cause of death^28^. To avoid chronic immunosuppression, research to evade immune rejection is very active. Multiple strategies, such as the generation of hypoimmune cells and targeted immunomodulation, have shown positive results in evading immune rejection^55–63^. The transplantation of hypoimmune cells without encapsulation and in a site without the possibility of retrieval requires SC-islets to be clearly defined to avoid irreversible complications.

Here, we applied two enrichment strategies for SC-α- and β-cells to generate defined 2^nd^ generation SC-islets. The first strategy utilised an iPSC fluorescent reporter line-based sorting to test the impact of endocrine cell-type composition on functionality *in vitro*. Next, we established cell surface antibody-based sorting methods to avoid the need for fluorescent-tagged cell lines and for upscaled production of SC-islets and *in vivo* assessment of functionality and safety. These strategies improved insulin secretion compared to non-enriched SC-islets. Transplantation experiments showed that changes in composition did not affect *in vivo* functionality, but that the percentage of off-target cells greatly influenced graft development. Moreover, our single cell transcriptomic analyses uncovered SC-islet cell-type composition before and after transplantation, identifying *in vitro* cell types responsible for unwanted growth *in vivo*. Altogether, our results showed that the generation of endocrine-enriched SC-islets does not greatly influence SC-islet function, but that enrichment reduces proliferative off-target cells and improves graft purity and safety long-term.

## Results

### Enrichment of SC-β-cells with SC-α-cells using flow sorting improves SC-islet functionality

To assess whether the functionality of SC-islets changes with different ratios of SC-α- and β-cells, we utilised the double reporter ARX^CFP/CFP^ x C-PEP^mCherry/+^ human iPSC line^64^. The reporter expression allows purification of hormone-producing SC-β- (INS^+^/C-PEP^+^), SC-α- (ARX^+^/GCG^+^), SC-α- progenitor- (ARX^+^/GCG^−^) and polyhormonal cells (ARX^+^/C-PEP^+^) using fluorescence-activated cell sorting (FACS). SC-islets were generated following an adopted three-dimensional (3D) differentiation protocol^65^, containing six stages (S1-6) with varied duration in each stage measured in days **(Fig. S1a)**. At the SC-islet stage, 3D aggregates were dissociated, sorted and engineered into 2^nd^ generation SC-islets containing defined ratios of SC-α- and β-cells **(Fig. 1a)**.

**Figure 1.**
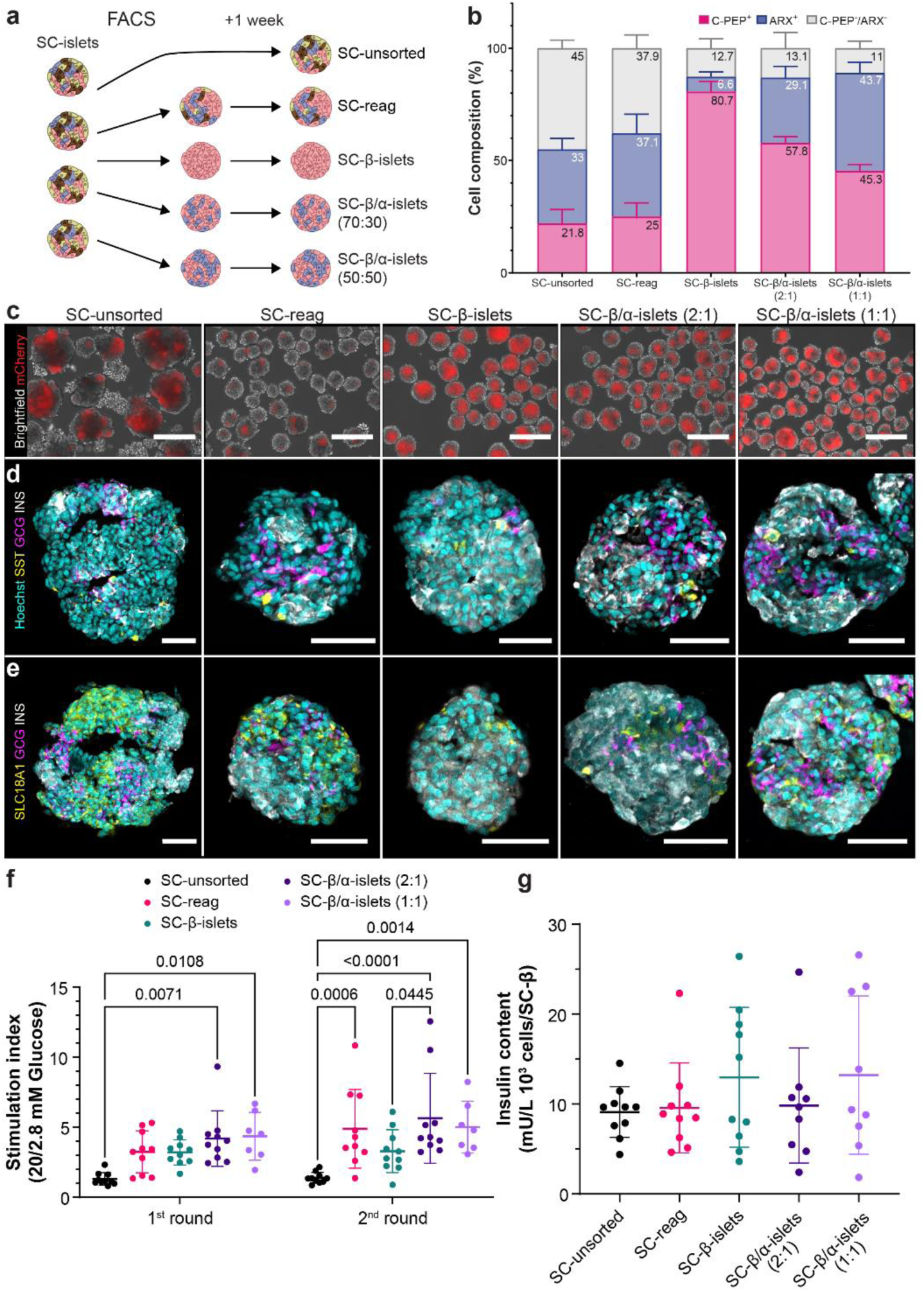
2^nd^ generation SC-islets with defined composition of SC-β- and α-cells showed improved functionality. (a) Experimental scheme to generate differently composed SC-islets. SC-unsorted islets are not dissociated at any time point of the differentiation. SC-islets were dissociated at S6.14 (Stage 6 day 14) and either reaggregated in Elplasia plates (SC-reag) or sorted by FACS for C-PEP^+^ and ARX^+^ and seeded in Elplasia at defined ratios. (b) Flow cytometric analysis of the cell composition using live cells (n=4). (c) Brightfield images of SC-islets with C-PEP (mCherry) expression 1 week after sorting. Scale bar, 300 µm. (d) Expression of islet cell-specific markers (SC-α- (GCG), SC-β- (INS) and SC-δ-cell (SST)) and (e) SC-EC-cell marker (SLC18A1). Scale bar, 50 µm. (f) Stimulation index of insulin secretion in response to high (20 mM) in comparison to low (2.8 mM) glucose (n=10, Two-way ANOVA statistical test with Tukey’s multiple comparison was used) (g) Insulin content determined by ELISA and normalised to 10^3^ cells and number of SC-β-cells (n=10, One-way ANOVA with Tukey’s multiple comparison).

We compared five types of SC-islets to assess whether different SC-islet composition impacts functionality: Two non-enriched control SC-islets and three SC-islets enriched for various SC-β/α-cell ratios. Control groups consisted of non-reaggregated (SC-unsorted, without dissociation) and a reaggregated benchmark control (SC-reag, dissociated and reaggregated without enrichment, as patented previously by Vertex Pharmaceuticals Inc)^65,66^. Enriched groups consisted of SC-β-islets (enriched for C-PEP/mCherry^+^), and SC-β/α-islets at 2:1 and 1:1 ratio (seeding at 70:30 and 50:50, respectively; C-PEP/mCherry^+^: ARX/CFP^+^). Purity of the FACS process varied depending on differentiation efficiency, with an average of 82.3 ± 5.3% for mCherry^+^ and 82.6 ± 6.1% for CFP^+^ **(Fig. S1b)**.

After 1 week in culture, the ratio of SC-β- and α-cells was maintained compared to the day of sort **(Fig. 1b),** and SC-β-cells were observable live through their mCherry reporter expression **(Fig. 1c)**. Reaggregation of SC-islets reduced the size and generated more homogenous clusters as expected^62,63^, but did not influence the SC-β- and α-cell composition when compared to unsorted SC-islets (compare SC-unsorted vs SC-reag in **Fig. 1b**). Further immunohistochemistry (IHC) analysis of cell-type composition reveals that all SC-islets contained SST^+^ SC-δ- **(Fig. 1d)** and SLC18A1^+^ SC-enterochromaffin-like (SC-EC) cells, an endocrine byproduct of the differentiation process **(Fig. 1e)**^39,41^. These results further indicate that a proportion of the C-PEP^−^/ARX^−^ cells consisted of other endocrine cells. Enrichment of SC-β- and α-cells resulted in visible depletion of SC-EC-cells **(Fig. 1e)**.

Functionality of SC-β-cells was assessed by insulin secretion in response to sequential low and high glucose stimulation **(Fig. S1c)**. Enriched SC-islets showed a higher total insulin secretion than non-enriched SC-islets upon glucose challenge. This test further revealed that reaggregation of SC-islets improved glucose response compared to SC-unsorted with similar SC-β- and α-cells proportion **(Fig. 1f)**. Within the group of enriched SC-islets, SC-β/α-islets (2:1) showed a significant higher stimulation index after repeated glucose challenge compared with SC-β-islets, suggesting that the presence and right ratio of SC-α-cells is beneficial for SC-β-cells functionality. A glucose ramp with an increase followed with a smaller decrease in glucose concentration further confirmed the increased glucose sensitivity and insulin secretion in all enriched SC-islet groups **(Fig. S1d)**.

The abundance of insulin protein in the different SC-islets reflected the increased number of SC-β-cells as shown before by cell-type composition **(Fig. S1e, 1b)**. While insulin content per SC-β-cell did not show a statistically significant increase after enrichment, a positive trend was observed **(Fig. 1g)**, suggesting that the improvement of glucose response may involve mechanisms beyond an elevated amount of insulin per cell. Taken together, we have established a reporter line-based sorting strategy to generate SC-islets with defined SC-β- and α-cell composition, reduced number of C-PEP^−^/ARX^−^ cells, and demonstrated an improved insulin secretion with SC-β/α-islets (2:1) in comparison to SC-unsorted and SC-β-islets.

### Cell surface marker-based enrichment upscales the generation of SC-islets with defined composition

The main limitation of a reporter-based enrichment through FACS is the low yield **(Fig. S1f)** and the need for genetically modified fluorescent PSC line. To enable upscaled production of enriched SC-islets for clinical applications, we explored cell surface antibodies suitable for magnetic activated cell sorting (MACS) to enrich for SC-β- and α-cells **(Fig. S2a)**. We initially tested several antibodies previously reported to mark endocrine cells. HPi1, a human pan-islet marker^67^, and the zinc dye TSQ, as a SC-β-cells marker^68–72^, did not demonstrate sufficient specificity to SC-β- or α-cells under our experimental conditions, which may reflect technical variability or limitations in their ability to distinguish the target cells **(Fig. S2b-c)**. Markers of a subset of primary human adult β-cells (NTPDase3)^48,73^ and α-cells (TM4SF4)^74^ were tested on SC-islets, however, no overlap was observed for the intended cell targets in SC-islets at S6 **(Fig. S2d-e)**. These antibodies have been tested for human islets and marked the respective populations; thus, technical issues can be ruled out (data not shown).

Integrin Alpha 1 (ITGA1) or CD49a is shown to reliably mark SC-β cells^39,50,75^, although it only labels a subpopulation of SC-β cells **(Fig. S2f)**. CD26 has been reported to mark adult human α-cells^76,77^. In SC-islets, CD26 recognised all SC-α-cells and some GCG^−^ cells **(Fig. S2g)**. Based on these results, we focused on CD49a and CD26 surface antibodies as these enabled reliable sorting **(Fig. S2h-i)** with high yield compared to FACS-based sorting **(Fig. S3a)**. MACS enrichment of SC-islets with CD49a resulted in 62.0 ± 5.8% mCherry^+^ and 17.9 ± 3.7% CFP^+^, and CD26 enrichment resulted in 4.9 ± 2.0% mCherry^+^ and 86.4 ± 3.3% CFP^+^ in subsequent FACS quantification **(Fig. S3b)**. After 1 week in culture, the efficient CD49a and CD26 enrichment for SC-β-cells and SC-α-cells, respectively, was maintained **(Fig. 2a)**. With both strategies, minor carryover of other endocrine cells, such as SC-δ- and EC-cells, was observed **(Fig. 2b)**.

**Figure 2.**
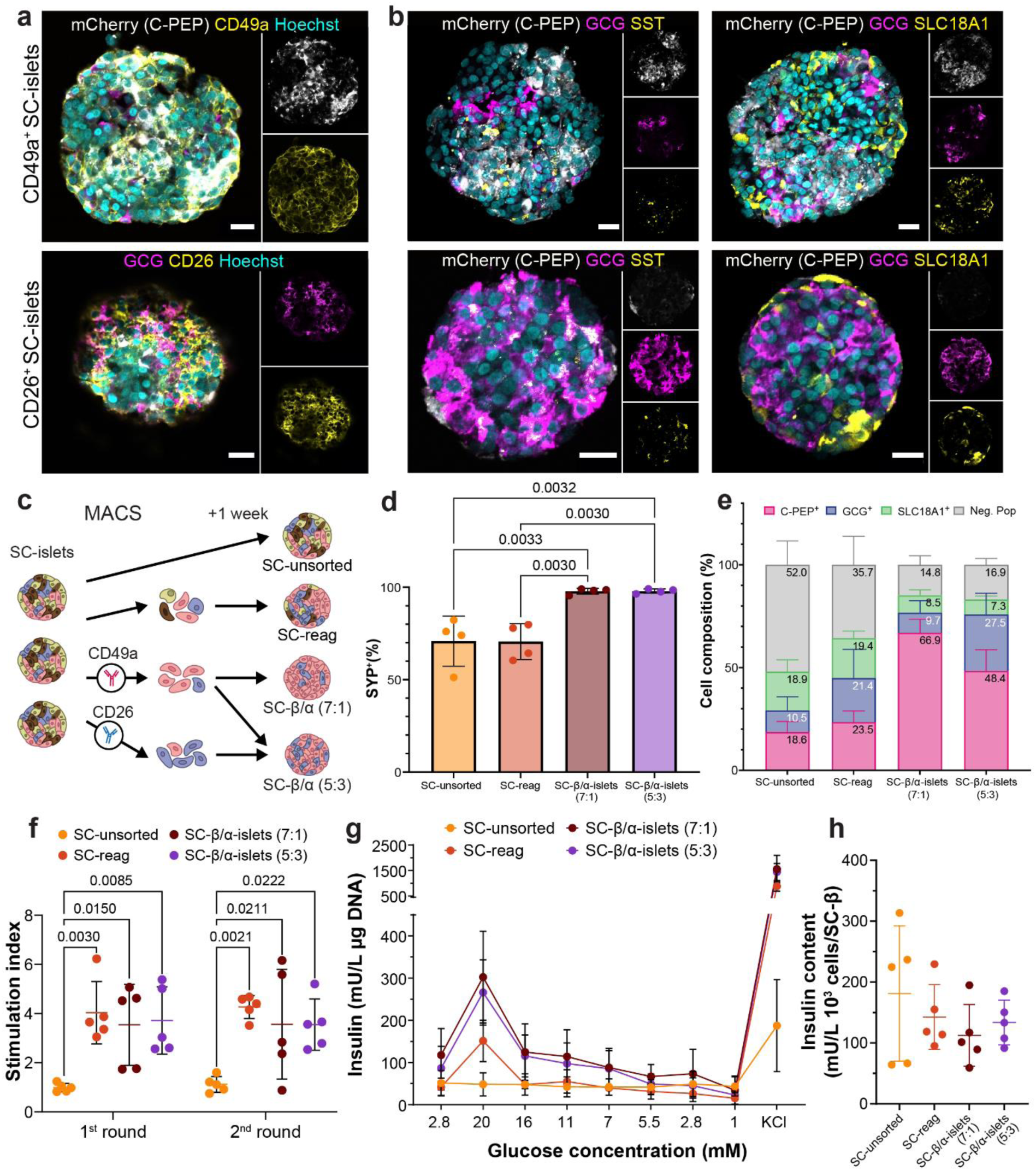
Antibody-based enrichment of SC-β- and α-cells generated defined endocrine-rich SC-islets. (a-b) Expression of respective antibody used for enrichment with endocrine cell markers for SC-α- (Glucagon), SC-β- (Insulin) and SC-δ- (Somatostatin) and SC-EC-cells (SLC18A1). Scale bar: 20 µm. (c) Experimental scheme for antibody-based sorting of SC-islets at S6.14. SC-β/α stands for SC-β/α-islets. (d) Flow cytometric quantification of pan-endocrine cells (Synaptophysin^+^,n=4, One-way ANOVA with Tukey’s multiple comparison) and (e) composition of SC-β-, SC-α-, and SC-EC-cells in enriched and non-enriched SC-islets. (f) Stimulation index of insulin secretion in response to high (20 mM) in comparison to low (2.8 mM) glucose (n=5, Two-way ANOVA statistical test with Tukey’s multiple comparison). (g) Insulin secretion in a glucose ramp (n=5). (h) Insulin content determined by ELISA and normalised to 10^3^ cells and number of SC-β-cells (n=5, One-way ANOVA with Tukey’s multiple comparison).

We used the combination of CD49a and CD26 to create SC-islets with defined ratios. We differentiated human iPSC without reporter (HMGUi001-A) to SC-islet stage, dissociated, MACS sorted and reaggregated the CD49a^+^ cells only, or CD49a^+^:CD26^+^ cells in 70:30 ratio **(Fig. 2c, S3c)**. Both enrichment strategies resulted in SC-islets containing >95% synaptophysin (SYP^+^) endocrine cells **(Fig. 2d)**. CD49a enrichment generated SC-islets containing ∼67% SC-β- and ∼10% α-cells, hereafter called SC-β/α-islets (7:1) **(Fig. 2e)**. Meanwhile the combination of CD49a and CD26 MACS generated SC-islets containing ∼48% SC-β- and ∼28% α-cells, subsequently called SC-β/α-islets (5:3) **(Fig. 2e)**. Presence of SC-δ-cells was observed in all enrichment strategies (**Fig. S3d**). Both sorting and reaggregation strategies also decreased the proportion of SC-EC-cells, from ∼19% to ∼9% in the 2^nd^ generation SC-islets **(Fig. 2e, S3e)**.

Functionality of these SC-islets was assessed by glucose-stimulated insulin secretion (GSIS). Enriched SC-islets showed higher total insulin secretion compared to non-enriched SC-islets **(Fig. S3f)**. SC-reag and enriched SC-islets showed a similar level of insulin response to glucose challenge compared to SC-unsorted **(Fig. 2f)**. In a similar fashion, SC-reag, SC-β/α-islets (7:1) and (5:3) exhibited a higher sensitivity to escalating and decreasing glucose concentration that was not observed in SC-unsorted **(Fig. 2g, S3g)**. The insulin content reflected the number of SC-β-cells **(Fig. S3h)**. However, no difference in insulin content per β-cell was observed **(Fig. 2h)**. This data shows that enrichment of SC-β- and α-cells and/or the reaggregation process improves insulin secretion upon glucose stimulation regardless of its endocrine composition *in vitro*.

### Cell surface marker-based sorting generated SC-islets with homogeneous transcriptional identity

To explore whether the endocrine enrichment alters the identity of the endocrine cells within SC-islets, we performed single-cell RNA sequencing (scRNA-seq) on SC-reag, SC-β/α-islets (7:1) and (5:3) one week after sorting. SC-islets were dispersed to a single-cell state and sorted for viability using FACS. Sorted live cells were then taken for further processing for scRNA-seq.

Single-cell transcriptomic analysis captured the main endocrine cell types shown in Uniform Manifold Approximation and Projection (UMAP) for dimensionality reduction, mainly SC-β-as marked by *INS* mRNA expression, α- (*GCG*), and EC-cells (*TPH1*; **Fig. 3a-d**), confirming our IHC and flow cytometry results **(Fig. 2e)**. Within the endocrine cells, a cluster of *ARX^+^* and *ARX^+^/CCK^+^* cells, as well as cycling cells belonging to SC-β-, α-, and EC-cell types, were identified. We further identified the presence of SC-δ- and ε-cells at a low proportion **(Fig. 3d)**. Consistent with our previous observation, CD49a sorting led to an enrichment of SC-β-cells subpopulation with high *INS* expression as seen by comparing SC-reag and SC-β/α-islets (7:1) in density plots **(Fig. 3e)** and single-cell *INS* expression **(Fig. S4d)**.

**Figure 3.**
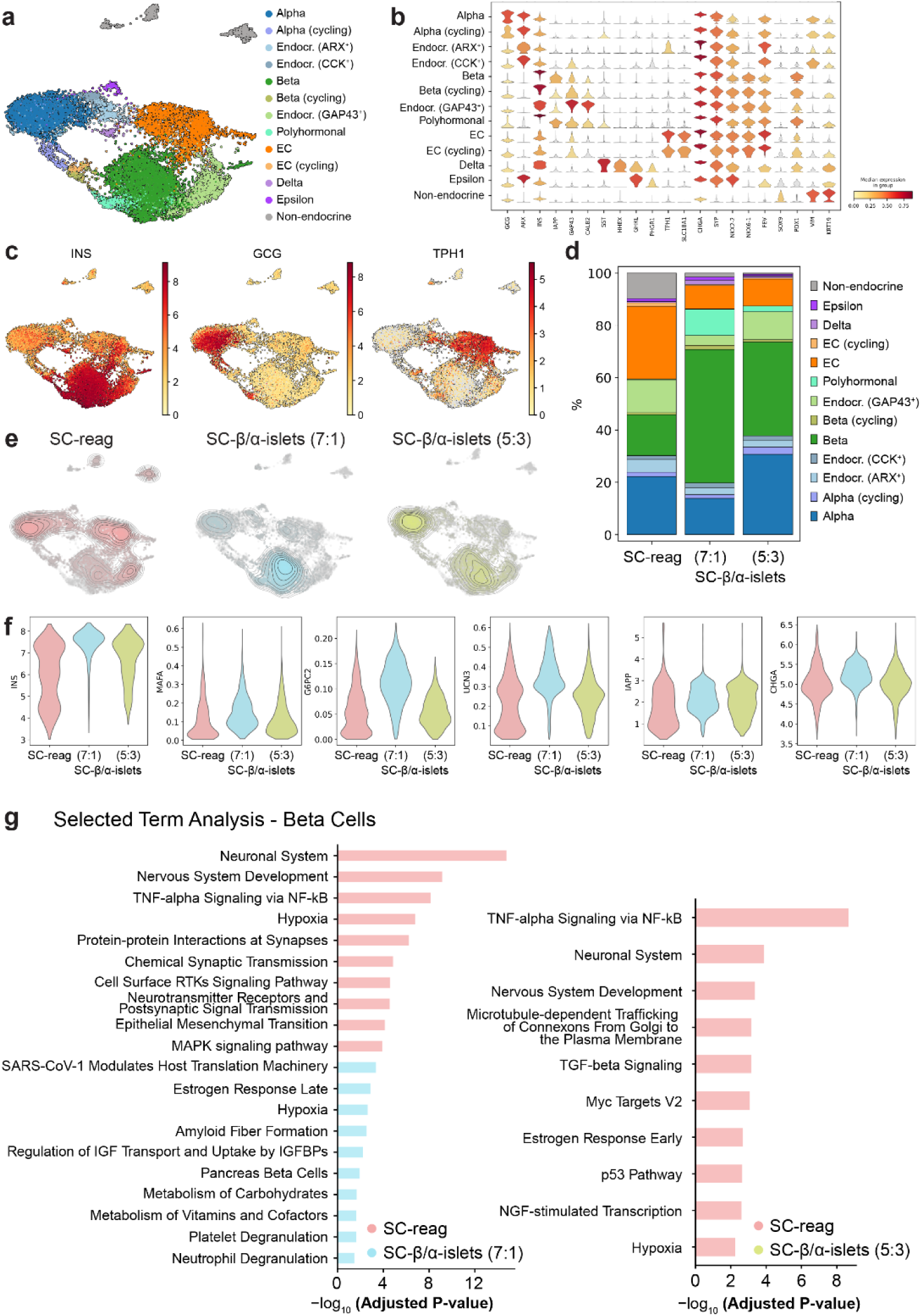
Transcriptional identity of SC-β-cells within different enrichment strategies exhibited minimal variation. (a) Cell type annotation shown in integrated Uniform Manifold Approximation and Projection for dimension reduction (UMAP) of in vitro samples. (b) Scaled log-normalised expression of marker genes for all clusters in this dataset. (c) UMAP of INS, GCG, and TPH1 expression. (d) scRNA-seq-based SC-islet composition on all cells. (e) Kernel density estimate plots showing the distribution of samples within the integrated UMAP. (f) Violin plots of maturation marker expression in SC-β-cells. (g) Selected Terms of the Over Representation Analysis in SC-β-cells between SC-reag and enriched SC-islets.

The SC-β-cells of SC-β/α-islets (7:1) showed a higher and more homogenous expression of *INS, MAFA, G6PC2, UCN3, IAPP and CHGA* compared to SC-reag **(Fig. 3f)**. This suggests that CD49a-enriched SC-islets show a more mature β-cell phenotype. β-cells specific changes in the expression of disallowed genes and genes associated with exocytosis, glycolysis, proliferation and islet signalling^20,78^ were similar between the groups **(Fig. S4e)**.

Further differential gene expression analysis (DGEA) between enrichment strategies mostly showed minimal differences as indicated by small log_2_ fold changes **(Fig. S5)**. The subsequent term enrichment analysis revealed an upregulation of TNF-α signalling via NF-kB, hypoxia, and apoptosis in SC-reag compared to enriched SC-islets **(Fig. 3g)**. Additionally, SC-β-cells of SC-reag upregulated neuronal processes that could indicate a more neuro/endocrine signature, while SC-β-cells of SC-β/α-islets (7:1) upregulated pathways, such as pancreas β-cells and metabolism of carbohydrates, fitting to a more mature β-cells phenotype. Interestingly, there were no significantly upregulated terms in SC-β/α-islets (5:3) compared with SC-reag, suggesting higher similarities **(Fig. 3g)**. This further indicates that the sorting combination of CD49a and CD26, enriches more SC-β-cell subpopulations **(Fig. S4d)**. In conclusion, manipulating SC-β- and α-cell composition in the SC-islets showed slight differences on SC-β-cells gene expression one week after sorting.

### SC-islets maintain normoglycemia independent of their cell-type composition

To assess *in vivo* functionality and safety of the SC-reag, as well as (7:1) and (5:3) enriched SC-islets, we transplanted these under the kidney capsule of non-diabetic immunodeficient mice and observed the graft function over 7 months **(Fig. 4a)**. Mouse β-cells were ablated through a single streptozotocin (STZ; 130 mg/kg) injection at month 3, after SC-islets had functionally engrafted to assess human β-cell function independently of the endogenous mouse β-cells^20^.

**Figure 4.**
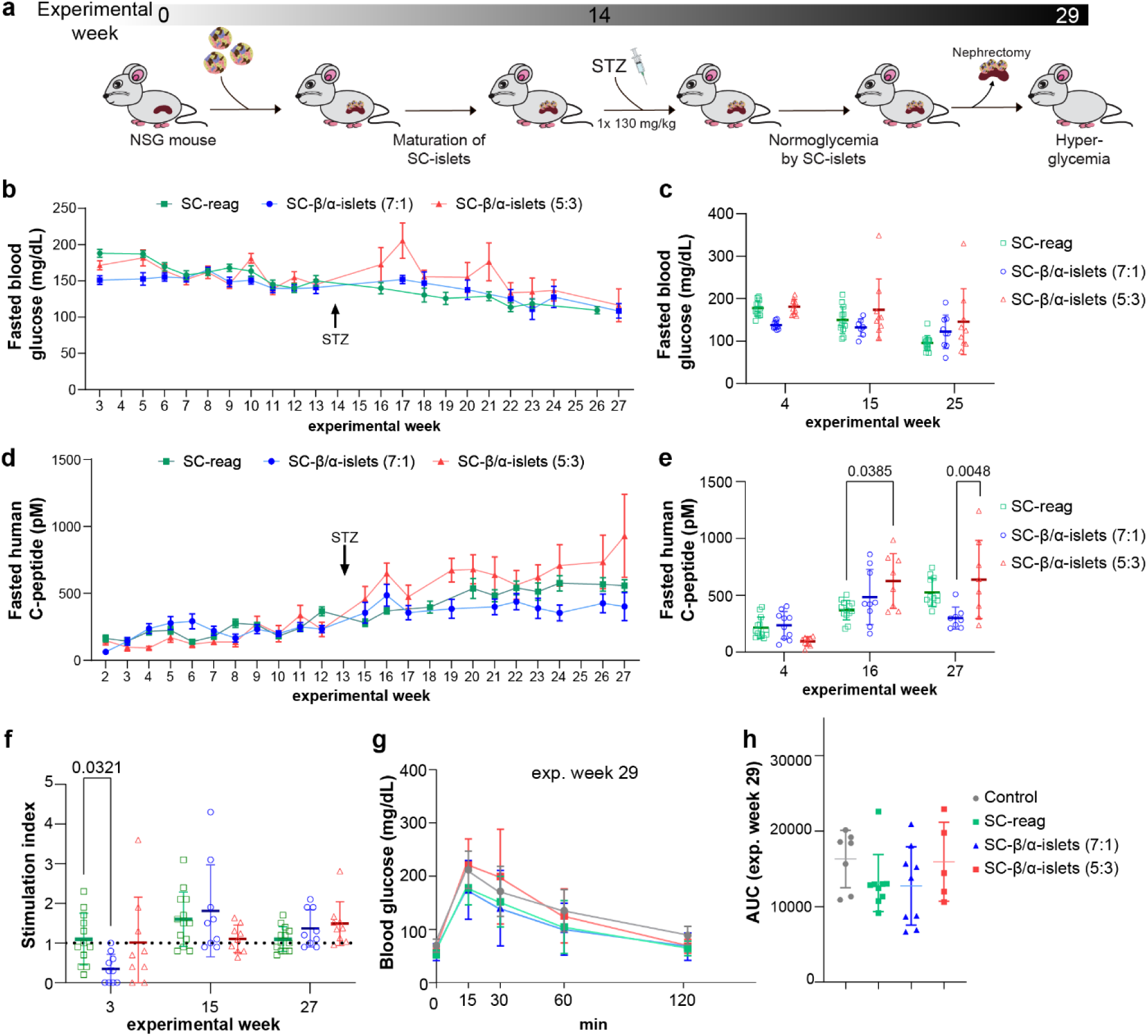
SC-islets maintained glycaemic control and improved in secretory capacity over 7 months in vivo. (a) Experimental workflow. (b) Blood glucose measurements after overnight 6 h fasting of SC-reag, SC-β/α-islets (7:1) and (5:3) over the experimental period (mean with SEM) and (c) at experimental start, after STZ injection, and experimental end (Tukey’s multiple comparison). (d) Human C-PEP measurements after overnight 6 h fasting of SC-reag, SC-β/α-islets (7:1) and (5:3) over the experimental period (SC-reag: n= 13, SC-β/α-islets (7:1): n=9-10, (5:3): n=9-10; mean with SEM) and (e) at experimental start, after STZ injection and experimental end (Šidák’s Test). (f) Stimulation index of glucose-stimulated insulin secretion after 60 min of glucose i.p injection (Tukey’s multiple comparison). (g) Glucose tolerance test (intraperitoneal) after 10 hours of fasting at experimental week 29 including non-transplanted, non-STZ injected control mice with (h) area under the curve (Kruskal-Wallis test with Dunn’s multiple comparison).

Fasted blood glucose decreases over time towards the human glycaemic set point^31^ **(Fig. 4b)** and comparison of glucose levels at the first and last time point as well as shortly after STZ injection revealed no significant difference **(Fig. 4c)**. Human C-peptide secretion after 6 hours of fasting increased over 7 months significantly in all three groups, indicating increase in graft function in follow-up and reliance of mouse blood glucose homeostasis on the engrafted human β-cells^20,79^. SC-β/α-islets (5:3) secreted the highest levels in comparison to the other two groups **(Fig. 4d-e)**. Notably, secretion levels in all three groups remained above the threshold required to maintain normoglycemia (>100 pM)^12^ following the destruction of endogenous β-cells. A significant increase in blood glucose levels was observed following kidney removal, further confirming the reliance of blood glucose regulation on the human grafts (**Fig. S4f)**.

Functionality of these SC-islets was assessed through *in vivo* GSIS and glucose-clearing assays **(Fig. 4f-h)**. At the beginning of the experimental period, SC-β/α-islets (7:1) showed a lower insulin response compared to SC-reag **(Fig. 4f)**. This was reversed at the end, with SC-β/α-islets (7:1) showing a slightly higher stimulation index, however, no significant differences were observed between the three transplanted SC-islet groups. Similarly, glucose clearing assays showed comparable functionality within the 3 groups, with no significant difference in the area under the curve **(Fig. 4g-h)**. Taken together, the tested SC-islet cell compositions did not significantly improve glycaemic regulation.

### Endocrine cells exhibit a mature phenotype after *in vivo* transplantation

To observe single-cell molecular signature differences between SC-islets before and after engraftment, histological and scRNA-seq analyses were performed on the explanted grafts after 7 months *in vivo*. Immunostaining on sectioned grafts showed continued presence of SC-β-, α-, EC-, and δ-cells in all SC-islets **(Fig. 5a)**, which is confirmed in our transcriptomic analysis **(Fig. 5b, S6a)**. Due to processing difficulties, we excluded SC-β/α-islets (5:3) after transplantation and integrated the scRNA-seq data from before and after transplantation (pre- and post-TX) to create a full overview of the changes that occur after SC-islet transplantation, engraftment and confirmed SC-islet function.

**Figure 5.**
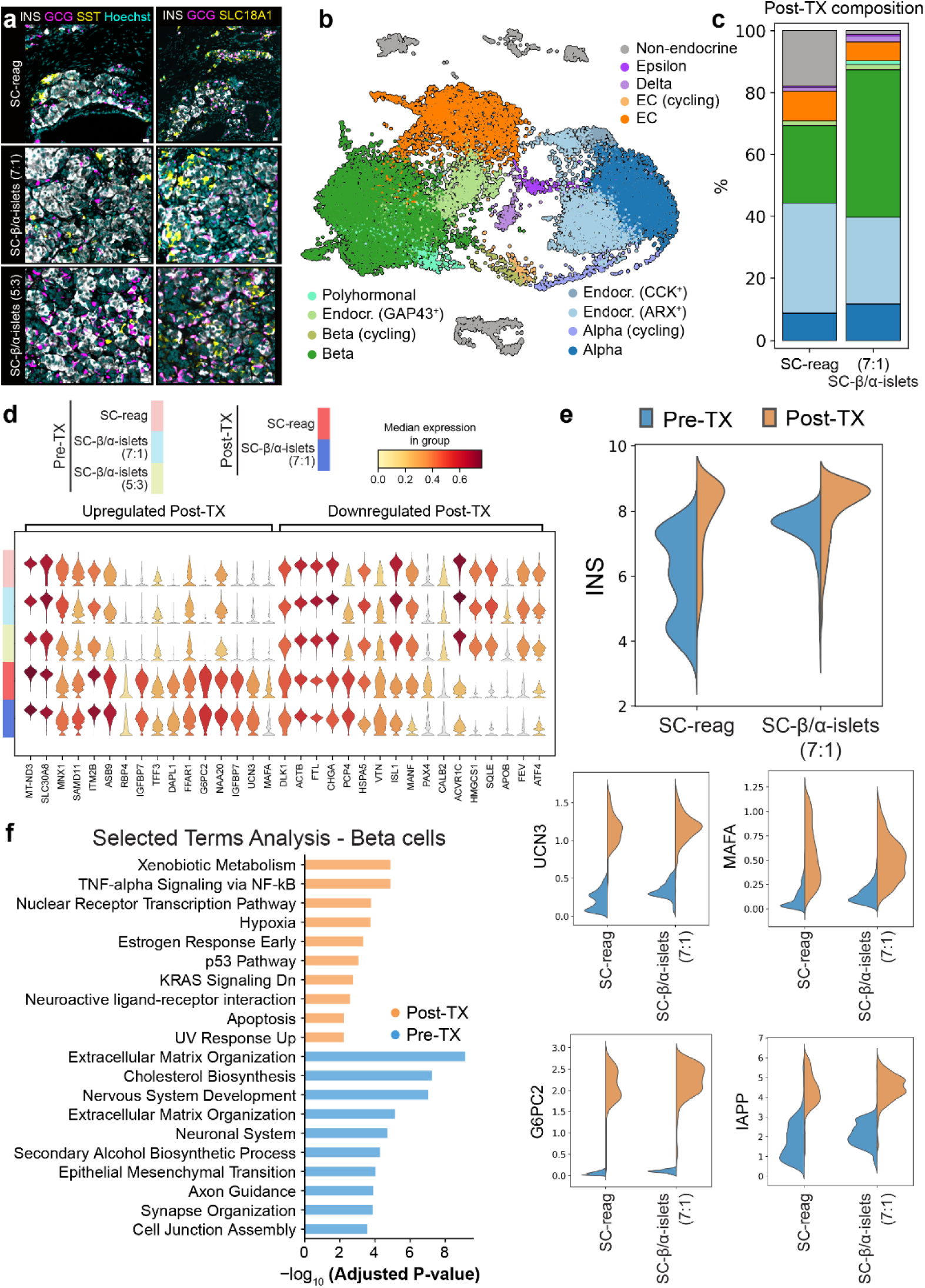
SC-β-cells showed more mature identity post-transplantation (post-TX). (a) IHC showing main endocrine cell types within SC-islets (Insulin – SC-β, Glucagon – SC-α, Somatostatin – SC-δ, SLC18A1 – SC-EC). Scale bar, 50 µm. (b) Integrated UMAP of all samples before (pre-) and after transplantation (post-TX) showing the cell type annotation. (c) Compositional bar plot depicting cell-type composition in post-TX samples. (d) Stacked violin plot (scaled log-normalised) of known genes to change after in vivo maturation. (e) β-cell maturation marker gene expression over the two timepoints between SC-reag and SC-β/α-islets (7:1). (f) Barplot of the selected pathways from the over-representation analysis of SC-β-cells before and after transplantation.

SC-reag showed an increase in the proportion of non-endocrine post-TX, meanwhile, no change was observed in SC-β/α-islets (7:1) **(Fig. 3d, 5c)**. Within the endocrine clusters of SC-reag and SC-β/α-islets (7:1), SC-β-cells appeared to be stable in proportion, while SC-α-cells were reduced. Interestingly, the amount of ARX^+^/GCG^−^ cluster was increased and the SC-EC-cells fraction was decreased post-engraftment in both SC-islets.

SC-β-cells of pre- and post-TX samples formed distinct clusters in the UMAP embedding, indicating the engraftment process influenced cell identity **(Fig. S6b-c)**. β-cell maturation after transplantation was further confirmed with gene expression changes **(Fig. 5d, S6d, S7b)**. Key β-cell identity and maturation genes (*INS, MAFA, G6PC2, UCN3, IAPP*) were upregulated after transplantation for both SC-reag and enriched SC-islets **(Fig. 5e, S7a)**. At post-TX, overall expression levels of the key β-cell genes between SC-reag and SC-β/α-islets (7:1) were comparable, however, SC-β/α-islets (7:1) exhibited a higher number of cells expressing genes at similar levels, suggesting reduced expression heterogeneity relative to SC-reag.

Pathways most changing in SC-β-cells post-TX included upregulation of genes related to TNF-α signalling via NF-kB, hypoxia, and xenobiotic response, while downregulation of pathways related to cholesterol biosynthesis and neuronal processes **(Fig. 5f)**. Changes were also observed in SC-α-cells with upregulation of *CHGB* and *LOXL4,* accompanied by lower expression of *ARX* **(Fig. S7c)**. Terms associated with circadian rhythm, estrogen response, and TNF-α signalling via NF-kB were upregulated in SC-α-cells post-TX while downregulating cholesterol homeostasis, epithelial-mesenchymal transition, and mTORC1 signalling **(Fig. S7d-e)**. Taken together, SC-β- and α-cells showed a more mature phenotype after transplantation.

### SC-reag off-target cells generated abnormal outgrowths compared to enriched SC-islets

A critical aspect in the generation of SC-islets, aside from functionality, is safety upon transplantation. To allow SC-islets for clinical use, it is vital to chart the *in vitro* cell populations responsible for potential adverse events.

As described, we characterised the on-target hormone-producing cells in different enrichment strategies. All SC-islets also contained a minor population of non-endocrine, SYP^−^ cells **(Fig. 2d)**. Seven months after transplantation, we observed an expansion of graft volume of SC-reag grafts compared to the day of transplantation **(Fig. 6a, S8a, S10a)**. Pathological analysis and pancreatic ductal marker CK19 staining revealed formation of cystic lesions of varied size and structure in addition to mesenchymal-like fibrosis, marked by PDGFRB **(Fig. 6b-c, S8b)**. This outgrowth did not occur in the grafts arising from enriched SC-islet preparations. While both enriched SC-islet preparations were devoid of large cysts and mesenchymal cells, they contained small CK19^+^ ductal regions, which were more common in grafts arising from SC-β/α-islets (5:3) in comparison to SC-β/α-islets (7:1) **(Fig. 6c, S8b)**.

**Figure 6.**
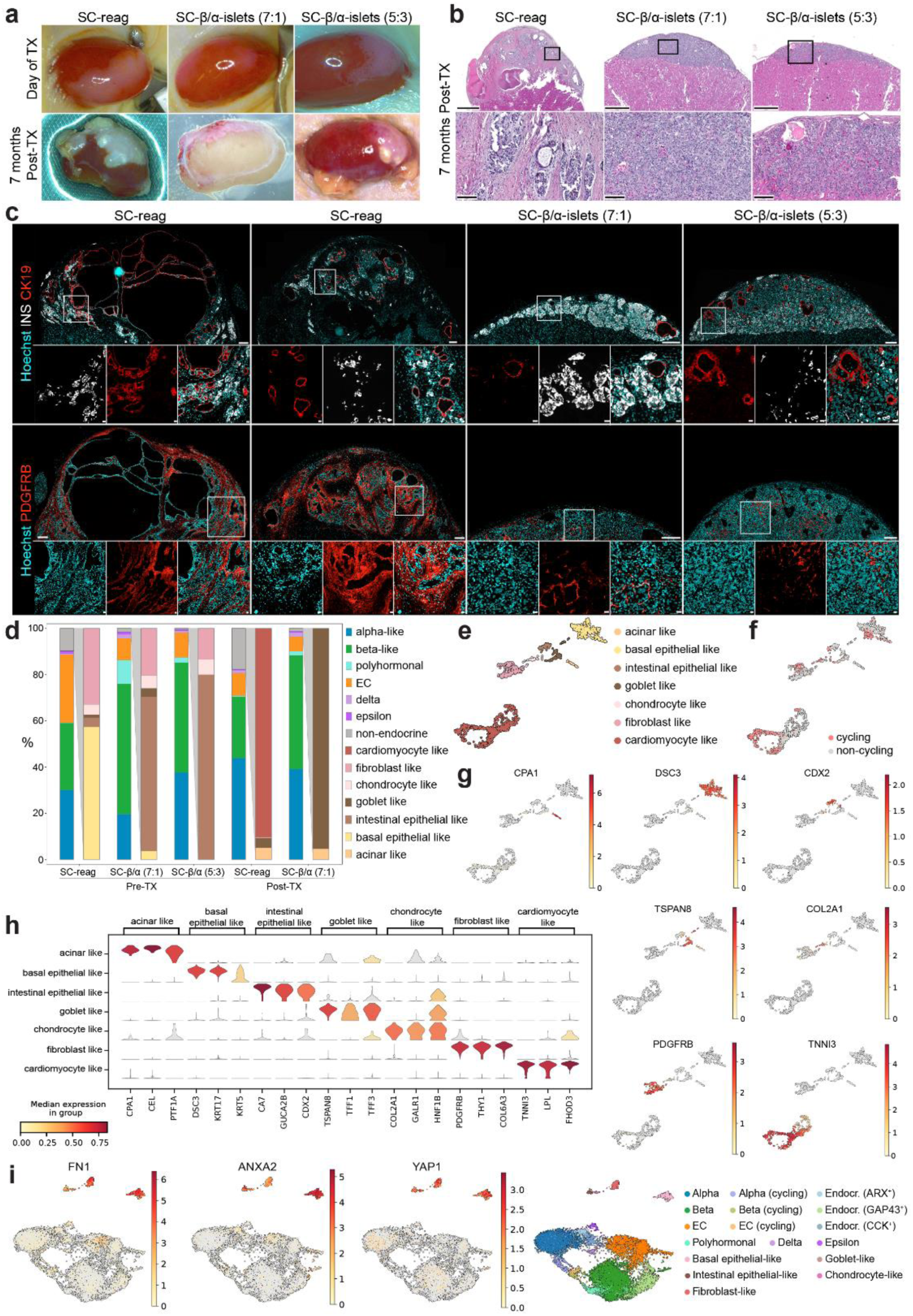
Enriched SC-islets contained minimal off-target cells resulting in pure grafts without outgrowths. (a) Images of graft on the day of transplantation (Day of TX) and after 7 months in vivo (7 months post-TX). (b) H&E staining of grafts 7 months post-TX. Scale bar, 500 µm and 100 µm. (c) IHC of paraffin-embedded grafts. Scale bar, 200 µm and 20 µm. (d) Composition plot of endocrine and off-target cells with closer distinction of off-target cells. SC-β/α stands for SC-β/α-islets. (e) UMAP-based embedding projection of off-target cells and (f) cycling cells within the off-target clusters. (g) UMAP-based embedding projection of distinct markers for each off-target cell cluster. (h) Expression of 3 distinct genes in each off-target cluster. (i) Expression of genes expressed exclusively in the non-endocrine cluster before transplantation.

Highly severe fibrosis marked with PDGFRB^+^ staining with invasion into the murine kidney was only observed in transplanted SC-reag islets **(Fig. 6b, 6c)**. Enriched SC-islet grafts showed mild fibrosis encapsulating the endocrine regions without infiltration into the kidney **(Fig. 6b)**. Additionally, osteoid metaplasia was found in SC-reag transplanted mice (12/13 mice) and SC-β/α-islets (5:3) grafts (3/9 mice; **Fig. S8c**), likely arising from multipotent mesenchymal stromal cells^80^. Notably, these bone-like structures were not observed in SC-β/α-islets (7:1) grafts (0/9 mice). These major differences are the result of the much higher proportion of non-endocrine cells in the reaggregated preparations (29.4% vs. 2.1% in SYP^−^ in histology **(Fig. 2d)** or 10% vs. 0.5-1.5% in scRNA-seq **(Fig. 3d)**).

To identify the source of the *in vivo* fibrosis, cystic lesions and outgrowths, we subclustered the non-endocrine cells in all transcriptomics samples **(Fig. 6d-e)**. The majority of the non-endocrine cells were found in both SC-reag pre- and post-TX samples **(Fig. 6d)**, indicating that the enrichment process was able to diminish the off-target cells. Interestingly, the proportions of non-endocrine cells varied within enrichment strategies. SC-reag contained more off-target cells with mesenchymal signature compared to SC-β/α-islets (7:1), which had more epithelial signatures. Within these cell types, we also identified off-target cells with proliferative capacity **(Fig. 6f).**

Based on their gene expression, we separated the off-target cells into seven subclusters **(Fig. 6g-h, S9)**. Two of them were of pancreatic lineage: an acinar-like cluster identified by *PTF1A* and *CPA1/2* expression, which was found only after transplantation, and a basal epithelial-like cluster, characterised by expression of *DSC3* and *KRT5*, which was found mainly in SC-reag pre-TX. We also identified two clusters that are likely of intestinal lineage. One with an enterocyte-signature marked mainly by *CDX2* and *CA7* found mostly before transplantation, while another with secretory-associated genes expressed in goblet-like cells marked with *TSPAN8* and *TFF3*, observed before and after transplantation. Interestingly, we observed the presence of CDX2^+^ cells post-TX, lining some cystic formations **(Fig. S10b)**.

The remaining three subclusters were marked by lower expression of the epithelial cell marker *EPCAM* and higher mesenchymal marker *VIM* expression **(Fig. S6c).** We detected chondrocyte-like cells marked with *COL2A1* expression and fibroblast-like cells expressing *PDGFRB* and *THY1*, which were mostly detected in all SC-islets pre-TX **(Fig. 6d, 6h)**. Immunostaining of PDGFRB uncovered that the proportion of fibroblast-like cells was present after transplantation and in higher proportion in SC-reag compared to the enriched SC-islets, which was not reflected in the composition of non-endocrine cells detected at the single-cell level **(Fig. 6c)**. A cluster of cardiomyocyte-like cells marked by *TNNI3 and FHOD3* expression was found exclusively in SC-reag after transplantation.

To develop markers suitable for use as release criteria, i.e. *in vitro* markers associated with *in vivo* outcome, we dissected gene expression of the *in vitro* off-target cells for markers absent from on-target cells. We identified *FN1*, *ANXA2*, and *YAP1* to be expressed exclusively in the non-endocrine cells, and their expression was detected in basal epithelial-like, intestinal epithelial-like, fibroblast-like, chondrocyte-like, and goblet-like clusters, covering all *in vitro* off-target clusters **(Fig. 6i)**.

## Discussion

Here, we tested the concept of engineering 2^nd^ generation SC-islets with increased safety and efficacy by defining the composition through enrichment strategies using reporter-based and cell surface antibody-based sorting strategies. Both resulted in almost pure endocrine-enriched SC-islets with defined SC-α- and β-cell compositions. Endocrine-enrichment resulted in improved SC-islet functionality *in vitro*, which did not directly translate to improved functionality *in vivo*. On the other hand, enrichment had a strong positive impact on graft purity and safety after transplantation. Based on scRNA-seq profiling, we described the origin of undesired off-target cells that arise during SC-islet differentiation and identified molecular markers for these cell populations. These off-target cells (ca. 10% non-endocrine cells in SC-reag, **Fig. 3d**) contribute to cystic lesions, mesenchymal-like fibrosis, osteoid metaplasia, as well as intestinal and exocrine outgrowths, compromising the safety of the grafted SC-islets. This highlights the importance of defining the final SC-islet product and formulate release criteria to ensure long-term safety, especially if a dose of 800 million cells per patient is required^28^ and hypoimmunogenic advanced therapy medicinal products (ATMPs) are anticipated to be transplanted in the future^63^.

Enrichment of endocrine cells increases overall insulin secretion of SC-islets, possibly due to the removal of inactive non-endocrine cells that inhibit intra-islet interconnectedness or simply due to better diffusion in the smaller SC-islet clusters^65^. The higher total insulin secretion would be beneficial for transplantation purposes, as fewer cells would be needed to achieve the effective does of the SC-islet product to achieve the intended insulin secretion level. This would achieve an economic advantage and would be easier for the manufacturing and distribution process, which is still one of the main challenges of β-cell replacement therapy^81^. Comparing enriched clusters, the presence of SC-α-cells is beneficial for SC-β-cell functionality *in vitro*, possibly through the known role of glucagon in stimulating insulin secretion^82^.

Using CD49a to sort for SC-β-cells and CD26 to sort for SC-α-cells increased the yield of endocrine cells. Specifically, CD49a enriches for a subpopulation of SC-β cells (ca. 50% of all generated SC-β-cells at S6.14) that is more mature on a transcriptomic level, as shown in our scRNA-seq analysis of *in vitro* samples **(Fig. 3f)**. However, this mature cell profile was not directly reflected in terms of SC-islet function *in vitro*. This might hint at the downside of β-cells subtype sorting using CD49a, as the result of antibody-based sorting was not on par compared to reporter-based sorting. It has recently been shown that both immature and mature β-cells contribute to islet operation^83^.

This could also mean that, comparatively, having SC-α-cells is more important than achieving an increase in SC-β-cell maturation for functionality. A recent publication showed that the presence of SC-α-cells and other islet cell types improves hypoglycaemia protection in diabetic mice, further cementing the importance of other cell types for glycaemia regulation^84^. Thus, sorting all SC-β- and α-cell subtypes to generate SC-islets with heterogeneous endocrine cells is the key to long-term functionality^34,72,83–88^. Additionally, CD26 was used in another setting to deplete a fraction of CD49a^+^ cells^50,75^. Despite being the best possible antibody combination reported to enrich for SC-α- and β-cells individually to obtain and characterise SC-islets with a defined ratio of endocrine cells, identification of specific (pan-) endocrine cell surface markers would be crucial for future research and to avoid the loss of 50% of the differentiated SC-β-cells after cell sorting.

Upon transplantation of non-enriched and enriched SC-islets, no significant functional differences between the groups were observed. This finding was accompanied by no significant differences in maturation profile between SC-β-cells of SC-β/α-islets (7:1) and SC-reag after transplantation, despite SC-β-cells of SC-β/α-islets (7:1) having a slightly more mature transcriptomic profile *in vitro*. This could be explained through continuous SC-β-cells maturation *in vivo*^20,78,79^. We speculated that during the 7 months *in vivo,* SC-β-cells of SC-reag islets continuously mature over time and eventually reach the maturity level of SC-β-cells of SC-β/α-islets (7:1).

Importantly, we have uncovered that enriched SC-islets showed no increase in size or outgrowth after 7 months *in vivo* thus reducing the potential safety risk. The only real off-target growth in our purified grafts was isolated cells of CK19^+^ cells, not organising in large cysts. A similar result was found in another study examining grafts with high SYP^+^ purity, with small cysts of CK19^+^ cells at a rate of around 1%^43^. SC-β/α-islets (5:3) derived grafts had more minor off-target outgrowths as opposed to SC-β/α-islets (7:1). This might be related to CD26 not being specific for only SC-α^77^, but possibly also expressed in ductal progenitors.

Using scRNA-seq profiling of SC-islets before and after transplantation, we observed that different starting *in vitro* off-target cells resulted in different *in vivo* outcomes **(Fig. S10c)**. We speculated that the fibrosis and osteoid metaplasia likely arose from the PDGFRB^+^ fibroblast-like and COL2A1^+^ chondrocyte-like cells^80^. These cell types possibly also give rise to the cardiomyocyte-like cluster. Meanwhile, off-target cells that are of intestinal lineage, intestinal epithelial-like and goblet-like cells, maintained their presence *in vivo*. Acinar-like cells likely emerged from basal epithelial-like cluster, further confirming the pancreatic nature of these cells. It is important to note that the identification of the subclusters was challenging, as most clusters showed a mix of cell markers and did not correspond to a human counterpart, suggesting that these cell types represent a dead end of *in vitro* differentiation. Additionally, as seen in the discrepancy between the amount of PDGFRB^+^ cells detected in IHC and scRNA-seq, it was not possible to digest the whole graft due to processing difficulties, which likely resulted into sampling biases.

We identified a set of marker genes expressed in all off-target clusters *in vitro* that could be used as future release criteria. Fibronectin, encoded by *FN1*, is an extracellular matrix (ECM) glycoprotein that is mostly expressed in stromal cells of the pancreas and is involved in cell-matrix adhesion, signalling and tissue repair^89–92^. *ANXA2* encodes for Annexin A2, a calcium-dependent phospholipid-binding protein that regulates cellular growth and is involved in signal transduction^93–96^. Finally, *YAP1* (Yes-Associated Protein) encodes a downstream effector of the Hippo signalling pathway that regulates cell proliferation, and its inhibition is essential for endocrine specification^97–99^. Interestingly, all three gene markers have a role in bone formation^100–106^.

Within the outgrowth, we did not see any indication of teratomas^107,108^ present in our grafts. This was supported by the lack of *OCT4* and *NANOG* detected in our scRNA-seq (data not shown). There is a possibility that other off-target cells might exist in different settings using other cell lines and/or differentiation protocols.

A previous study has established a method to expand unwanted outgrowths *in vitro* to mimic the potential outcome from transplantation^44^. In the study, they used EGF to stimulate the off-target growth. This is consistent with our scRNA-seq result, which uncovered that *EGFR* is expressed in all off-target clusters, including before and after transplantation **(Fig. S9b)**. This further highlights the importance of taking a complete look at the whole graft in scRNA-seq data and not exclusively focusing on the endocrine compartment.

In summary, here we have highlighted the importance of generating defined 2^nd^ generation SC-islets through positive selection of endocrine cells. This enrichment minimises the risk of off-target outgrowths *in vivo*. A combination of pathological analysis and scRNA-seq has enabled the identification of off-target markers. This identification is crucial to perform quick and reliable quality and release controls, to ensure long-term safety of the SC-islets upon transplantation, building on the recent advancements in the field of islet replacement therapy.

## Supporting information

Supplementary Table 4 - Differential Gene Expression

Supplementary Table 5 - Terms Analysis

## Resources availability

Source data and scripts for scRNA-seq analysis will be made available upon publication. Any other data supporting the findings of this study are available from the corresponding author on reasonable request.

## Acknowledgement

We are thankful to L. Appel, E.V. Baumgart, K. Diemer, F.J. Farkas, and I. Kunze for the technical support. Our sincere thanks go to C. Karampelias and C. Cozzitorto for the helpful comments and discussions. We are grateful to D.M. Thomson, M. Catani, and E. Schlüssel for the administrative support. We thank the Core Facility Laboratory Animal Services and the animal caretakers of Helmholtz Munich, particularly F. Schwarz and A. Brähler. We acknowledge the support of Core Facility Pathology and Tissue Analytics, especially A. Feuchtinger, M. Tost, and U. Buchholz for providing pathological analysis and the technical support of Genomics (CF-GEN) at Helmholtz Munich. We thank T. Walzthöni for help with secondary processing. This work has received funding from the European Union’s Horizon 2020 research and innovation programme under grant agreement ISLET number 874839. This work was further supported by BMBF with the project “eISLET” (grant number 031L0251) and DFG Research Infrastructure NGS_CC (project 407495230) as part of the Next Generation Sequencing Competence Network (project 423957469). NGS analyses were carried out at the Competence Centre for Genomic Analysis (Kiel). Further funding was provided by the Helmholtz-Gemeinschaft and German Center for Diabetes Research (DZD e.V.).

## Author contributions

E.S.A.S., N.K.R. and H.L. conceived and designed the study, and wrote the manuscript. K.S. and S.S. conceived and designed the study. E.S.A.S., N.K.R., K.S., S.S. and T.Ö. performed the experiments and analysed the data. H.R. and M.S. performed and analysed the scRNA-seq study. S.F. supported the scRNA-seq study. V.L. participated in data interpretation and manuscript writing. H.L. supervised the study, provided resources, and acquired the funding.

## Methods

### *In vitro* differentiation of hiPSCs towards SC-islets

Human induced pluripotent stem cell (hiPSC) HMGUi001-A^109^ and ARX^CFP/CFP^ x C-PEP^mCherry/+^ (HMGUi001-A-54)^64^ cell lines were used in this project.

hiPSCs were maintained in Geltrex^TM^ (Life Technologies, A1413302) coated plates with StemMACS^TM^ iPS-Brew XF medium (Miltenyi Biotec, 130-104-368) and passaged every 3-4 days at > 80% confluency. After adjustment in 2D format, cells were dissociated with Accutase® solution (Sigma-Aldrich, A6964), seeded in 30 ml spinner flasks (Reprocell, ABBWVS03A-6) at an 8 × 10^5^ cells/ml density, and maintained at 60 rpm. hiPSC aggregates were passaged every 3-4 days using Accutase® solution.

Differentiation towards SC-islets was started 3-4 days after passaging, as previously described^65^. In brief, aggregates in 3D were exposed to different media conditions that correspond to different stages of differentiation. Media composition and cytokines combination of the 6-stage differentiation protocol are listed in Supplementary Table 1-2.

### SC-α- and SC-β-cells sorting

SC-islets at S6.14 (stage 6 day 14) were dissociated into single cells using TrypLE^TM^ Select (ThermoFisher Scientific, 12563011) and strained through 40 µm cell strainer to remove clumps. A small portion of dissociated cells was kept without further processing and seeded immediately for SC-reag. For ARX^CFP/CFP^ x C-PEP^mCherry/+^ reporter line sorting using FACS, single cells were stained for live/dead using DAPI (Carl Roth, 6335.1) for 15 minutes and washed with PBS. Then, the cell pellet was resuspended in FACS Buffer (2% BSA and 0.2 mM EDTA in PBS) and run through FACS Aria^TM^ III (BD Biosciences). Sorted cells are collected, counted, and seeded in 96-well Elplasia® plates (Corning, 4442) in a seeding density of 10^5^ cells/well with S6 media supplemented with 10 µM ROCK inhibitor (S6+ROCKi, Y-27632, Biotechne, 1254/10). Two days after sorting, sorted SC-islets were moved to Costar^TM^ Ultra-Low Attachment plates (Fisher Scientific, 10154431), shaking at 90-100 rpm for another 5 days. The media was refreshed every 2 days.

For cell surface antibody sorting, single cells were counted and resuspended in FACS Buffer (12.5 µl/10^6^ cells). Antibodies were added to the cells: CD49a-PE and CD26-PE antibody, then incubated at 4°C for 20 minutes. Afterwards, cells were washed once with PBS to remove remaining antibodies followed by incubation with Anti-PE microbeads (1 µl in 10 µl FACS buffer/10^6^ cells) at 4°C for 15 minutes. Cells were washed once with PBS and resuspended in 500 µl FACS Buffer. Magnetic sorting was performed based on the manufacturer’s protocol. The enriched cells were then seeded at a density of 0.5 × 10^6^ cells/ml in a 6-well Ultra-Low Attachment plate and kept on a shaker at 90-100 rpm. Media was refreshed every 2 days, and SC-islets were used for analysis 7 days after sorting.

### Insulin secretion assay (*in vitro*)

Krebs Ringer buffer (KRBH; 129 mM NaCl, 4.8 mM KCl, 1.2 mM KH_2_PO_4_, 1.2 mM Mg_2_SO_4_-H_2_O, 2 mM CaCl_2_, 24 mM NaHCO_3_, 6 mM HEPES, 0.2% BSA in Millipore water pH 7.4) was prepared and used to prepare low glucose (LG, 2.8 mM glucose), high glucose (HG, 20 mM glucose) and KCl (25 mM KCl) solutions. Ten SC-islets (∼10,000 cells) were picked, washed two times with KRBH, and equilibrated with LG for at least 1.5 hour followed with HG for 30 minutes at 37°C. Then, SC-islets were incubated in a sequential series of LG1-HG1-LG2-HG2-KCl. SC-islets were exposed to the glucose solutions for 1 hour, and KCl solution for 30 minutes. Supernatants were collected after each round, and SC-islets were washed once after the HG rounds. At the end of the experiment, SC-islets were digested for DNA for normalisation. Insulin measurement was performed using Mercodia Human Insulin ELISA Kit (Mercodia, 10-1113-10) following the manufacturer’s protocol.

### Glucose ramp (*in vitro*)

Static glucose ramp was started by preparing sterile KRBH solution as previously described. Glucose solutions of 1, 2.8, 5.5, 7, 11, 16, 20 mM were prepared along with a 25 mM KCl solution. Ten SC-islets were picked, washed two times with KRBH, and equilibrated with 2.8 mM solution for 1 hour and 20 mM for 30 minutes at 37°C. Then, SC-islets were exposed to the different glucose concentrations in a sequential manner (2.8-20-16-11-7-5.5-2.8-1-KCl) for 30 minutes each. Supernatants were collected and SC-islets were washed with KRBH between each step. Insulin was measured using Mercodia Human Insulin ELISA Kit following the manufacturer’s protocol and normalised by DNA content.

### DNA Isolation

SC-islets were lysed with DNA lysis buffer and treated with Proteinase K (Roche, 3508838103) overnight at 60°C. DNA was precipitated with NaCl and ethanol, washed three times with 70% ethanol and ultimately resuspended in TE buffer (ddH_2_O, 10 mM Tris pH 7.6, 1 mM EDTA).

### Insulin content

Insulin content of the SC-islets was extracted following the protocol as previously described^17^. Briefly, SC-islets were washed with PBS and dissociated using TrypLE Select. Cells were counted, and the dissociated cells were resuspended in Acid-Ethanol solution (1.5% HCl and 70% EtOH). Tubes were kept on a shaker at 4°C overnight, then centrifuged at 2,100*g* for 15 minutes. Supernatants were collected and neutralised with an equal volume of 1 M Tris-HCl (pH 7.4). Measured insulin was normalised by cell number (10^3^ cells).

### IHC staining

SC-islets were fixed with 4% paraformaldehyde (PFA) for 20 minutes. Samples were then exposed to a sucrose gradient for dehydration (10-30%), embedded in tissue freezing medium (OCT, Leica, 14020108926) and frozen for cryosection. SC-islets blocks were sectioned at a thickness of 12-15 µm. Sections were permeabilised for 30 minutes with Permeabilisation Buffer (0.1% Triton-X with 0.1M Glycine in PBS). The sections are blocked for at least 1 hour using blocking serum (10% FBS, 0.1% BSA, 3% donkey serum, 0.1% Tween-20 in PBS) and stained for the marker of interest overnight at 4°C. List of primary antibodies can be found in Table 3. Following overnight incubation, sections were stained with secondary antibody for at least 2 hours at room temperature. The nucleus was indicated with Hoechst (Life technologies,62249). Images were taken with a Zeiss LSM 880 Airy Scan confocal microscope and analysed using Zeiss Zen Blue software.

### Flow cytometry analysis

SC-islets were dissociated into single cells using TrypLE Select and fixed with 4% PFA for 10 minutes. Single cells were then permeabilised with Permeabilisation Buffer for 15 minutes and stained with antibodies. Incubation length for conjugated antibodies was at least 1 hour at room temperature. Unstained samples were taken as a control for staining. Stained cells were analysed using BD FACS Aria III. List of antibodies can be found in Table 3.

### Ethics and animal maintenance

Animal experiments were performed at the central facilities at HMGU following the German animal welfare legislation and acknowledged guidelines of the Society of Laboratory Animals (GV-SOLAS) and of the Federation of Laboratory Animal Science Associations (FELASA). For all experiments, male vNOD.Cg-Prkdcscid Il2rgtm1Wjl/SzJ mice were purchased from Charles River Laboratories. Mice used for transplantation were between the ages of 29 and 56 days and weighing >27 g. Mice were held at a 12h day/light cycle, fed with Altromin (1318 best.) and received acidified water (0.2mM HCl) ad libitum.

### Transplantation of SC-islets under the kidney capsule

ARX^CFP/CFP^ x C-PEP^mCherry/+^ hiPSC reporter line was used for all transplantation experiments. SC-islets were transplanted in aggregate form with amounts equal to 3 million cells/mouse. After ensuring surgical tolerance using the toe muscle reflex, mice were shaved on the side of the abdomen (above the kidney). An incision was made in the skin and the peritoneum to reveal the right kidney. A hole using a sharp forceps was made in the kidney capsule for the insertion of SC-islets using a Hamilton syringe (Neolab Migge GmbH, HU-1497). Images of engrafted SC-islets were taken with Leica S9i (Leica LED3000 RL, 58 mm). The wound was closed with suture, wound clip and wound glue. After 3 months, streptozotocin (STZ; Sigma-Aldrich, S0130), resuspended in Citrate buffer pH 4.5, was i.p. injected.

### Glucose-stimulated insulin secretion assay (*in vivo*)

Mice were day-fasted for 6 hours by whole cage change. Blood was sampled for human and murine C-PEP after fasting. Sample for human C-PEP was collected 60 min after i.p. glucose (2g/kg BW) injection. Glucose was diluted in saline solution and sterile filtered. Mice were held in a restrainer, after which the tail was disinfected and a 24G needle (B. Braun, TZ-1446) was briefly inserted into the lateral caudal tail vein. Murine blood was collected in EDTA-coated microvettes (SARSTEDT AG & Co. KG, 16444) and then centrifuged for 10 min at 4°C. Blood serum was collected in PCR tubes and frozen at −20°C. Human and murine C-PEP were measured using ELISA kits (Mercodia, 10-1136-01 and 90050) according to the manufacturer’s instructions. Absorbance was assessed using a Plate reader (Thermo Fischer Scientific, Varioscan Lux).

### Glucose tolerance test (ipGTT)

Mice were fasted for 10 hours at night by being placed in a new cage containing a plastic house and bedding. Blood glucose was measured at 0, 15, 30, 60 and 120 min after i.p. glucose (2 g/kg BW) injection. For blood glucose assessment, the tail tip was punctured with a 24G needle and measured with a glucometer (Contour next).

### Nephrectomy – Removal of the kidney with engrafted SC-islets

Mice were anaesthetised, analgesized, and kidney was exposed as stated before. Fat and fibrotic tissue were carefully pulled off the kidney before exposing the kidney from below the peritoneum. SILKAM thread (6/0, 45cm, DS 16, non-absorbable) was prepared in the form of a loop and placed under the kidney, around the renal pelvis and tightly knotted. The kidney was removed by using sharp scissors to cut underneath the knotted thread. The incisions were closed as previously stated.

### Kidney processing for IHC

Kidneys used for IHC were washed in -/-PBS and fixed with 4% (w/v) formalin (Sigma Aldrich, HT501128) at 4°C overnight for cryofreezing or paraffin. Samples for cryofreezing were dehydrated in a 6-well plate in 10 % Sucrose for 1 h at room temperature, 30% Sucrose (ITW Reagents, PanReac AppliChem, 57-50-1) overnight at 4°C and 30% Sucrose:OCT (1:1) at 4°C overnight. Organs were embedded in moulds (Sigma-Aldrich, E4140-1EA) containing OCT and stored at −80°C. Samples were thawed to −20 °C, cut 10-15 µm with a cryostat (Leica, CM1860) and placed on glass slides (Thermo Fisher Scientific, J1800AMNZ). Samples embedded in paraffin were cut into 3 µm slices for hematoxylin and eosin (H&E) and Sirius Red staining. The stained tissue sections were scanned with an AxioScan 7 digital slide scanner (Zeiss, Oberkochen, Germany) equipped with a 20x magnification objective.

### Sample processing for scRNA-seq

SC-islets were dissociated using TrypLE Select at 37°C until a single-cell suspension was reached. The reaction was stopped with S6+ROCKi and cells were stained for live/dead using DAPI at 4°C for 15 min. The stained cells were sorted for viability using FACS Aria^TM^ III and sorted live cells were taken for further processing.

Graft was surgically removed in -/- PBS with premoistened forceps and scissors and kept on ice until further processing. Transplants were dissociated using TrypLE Select supplemented with ROCKi at 37°C for 10 min or until a single-cell suspension was reached. The reaction was stopped with PBS with 1% BSA. Cells were washed with PBS containing 2% FCS and diluted to a final concentration of ∼1000 cells/µl in PBS containing 2% FCS. After quality control and counting, the cell suspension was immediately used for scRNA-seq library preparation with a target recovery of 10000 cells.

Libraries were prepared using the Chromium Single Cell 3ʹ Reagent Kits v3.1 (10x Genomics, 1000268) according to the manufacturer’s instructions. Libraries were pooled and sequenced on an Illumina NovaSeq6000 with a target read depth of 50000 reads/cell. FASTQ files were aligned to the GRCh38 human and GRCm39 mouse genomes to distinguish host from graft cells and pre-processed using the CellRanger software v7.1.0 (10x Genomics) for downstream analyses.

### scRNA-seq analysis

Analyses were performed with Scanpy (Python v3.12.4 or v3.10.13, scanpy v1.10.4 or 1.9.8, anndata v0.11.3 or 0.10.6)^110^, unless stated otherwise. All samples were filtered for a minimum of 1 count per cell, as well as a minimum of 1 cell per gene. Empty droplets and ambient gene probability were calculated using the emptyDrops (FDR ≤ 0.05) function of DropletUtils (R v4.3.1, v1.22.0)^111^. Cells with a mitochondrial fraction over 25%, counts less than 6,000 or higher than 125,000 and less than 4000 genes were excluded. In some samples, thresholds were set stricter, according to the respective sample’s quality. The ambient probability threshold was set at 0.0006.

Doublet detection was performed with scrublet (scanpy v1.9.8, sim_doublet_ratio=5, expected_doublet_rate=0.09, threshold=0.25)^112^, ScDS (R v4.2.2, v1.14.0)^113^, scDblFinder (R v4.2.2, v1.12.0)^114^, doubletdetection (n_iters=200, random_state=123, pval ≤10^−6^, v4.2)^115^, DoubletFinder (R v4.2.2, v 2.0.3)^116^ and solo (n_hidden=256, n_latent=20, gene_likelihood=’nb’, scvi v1.1.2)^117^. The samples were normalised and integrated into 3 groups: pre-TX, post-TX and all. Normalisation was performed with R (Seurat v5.1.0, R v4.4.2) packages scran (v1.30.2)^118^ and sctransform (v0.4.1)^119^.

The initial embedding for preliminary cell type annotation was calculated on the PCA space using the top 4000 variable genes. Clusters were identified with leiden^120^ and annotated with marker genes (*INS, GCG, SST, ARX, TPH1, MKI67, CHGA, KRT19, VIM, EPCAM, GAP43, RPS26, S100A11, TPT1, NDUFA4, COL1A2*). The samples were then integrated with scvi (v1.2.2, n_hidden=512, n_latent=50, n_layers=2) on the whole transcriptome with ‘sample’ as batch key and the stage as additional covariate in the pre- and post-TX integration.

Using the resulting UMAP and leiden clusters, aggregated doublet clusters and cells labelled as doublets with more than 3 of the detection methods were excluded. The remaining doublet was excluded, and the initially annotated object was then integrated using scANVI. The gene counts were denoised and imputed with scanpy’s magic implementation (scanpy v1.11.0)^121^ and a finer annotation was performed on the doublet excluded, integrated object using *EPCAM, CHGA, MKI67, FANCI, ARL6IP1, INS, GAP43, CALB2, NPTX2, ASPH, PCDH7, MAP2, GCG, ARX, APOH, DPP4, ONECUT3, GC, TTR, SPINK1, SST, GHRL, TPH1, SLC18A1, STAC, GLIS3, SLC30A8, ERO1B, DDIT3, IAPP, VGF, KRT19, VIM, GLIS3, ANXA1, BICC1, CPA2, PTF1A, REG1A, SOX9, FHOD3, LGALS1, COL1A2, TNNI3* as marker genes, either from prior knowledge or identified with scanpy.tl.rank_genes_groups.

Specific marker genes for the nonendocrine clusters were identified with sc2marker (R v 4.4.2, v1.0.3)^122^. Differentially expressed genes within cell types and among enrichment strategies or stages were identified with DElegate’s (R v4.4.2, v1.2.1)^123^ findDE method (method = “edgeR”). Log2FoldChange and p-value thresholds were set dynamically depending on the distribution, based on z-scores. The lists of the top differentially expressed genes for each comparison can be found in Supplementary Table 4.

Known β-cell maturation markers (*INS*, *G6PC2*, *UCN3*, *IAPP*, *CHGA* and *MAFA*) and α-cell markers were used to compare enrichment strategies (*ARX, GCG, LOXL4, CHGB*). Over Representation analysis (ORA) was performed with gseapy’s (v1.1.7)^124^ enrichr implementation with a padj cutoff of 0.05. The genesets used were KEGG (version 2021)^125^, HALLMARK (version 2020)^126^, GO biological processes (version 2025)^127,128^, and Reactome (version 2024)^129^. The combined results were either filtered by keywords or aggregated by jaccard index. The term with the highest combined score or the term present in the exact term list (Supplementary Table 5) was kept.

### Statistical analysis

All statistical analyses were performed using GraphPad Prism v10. Statistical analysis used depends on the sample, as described in the figure legends. Error bar indicated standard deviation, unless specified.

