## Supplementary Figures 1-10 and Supplementary Tables 1-3 for "Endocrine-enriched stem cell-derived islets improve long-term safety *in vivo*"

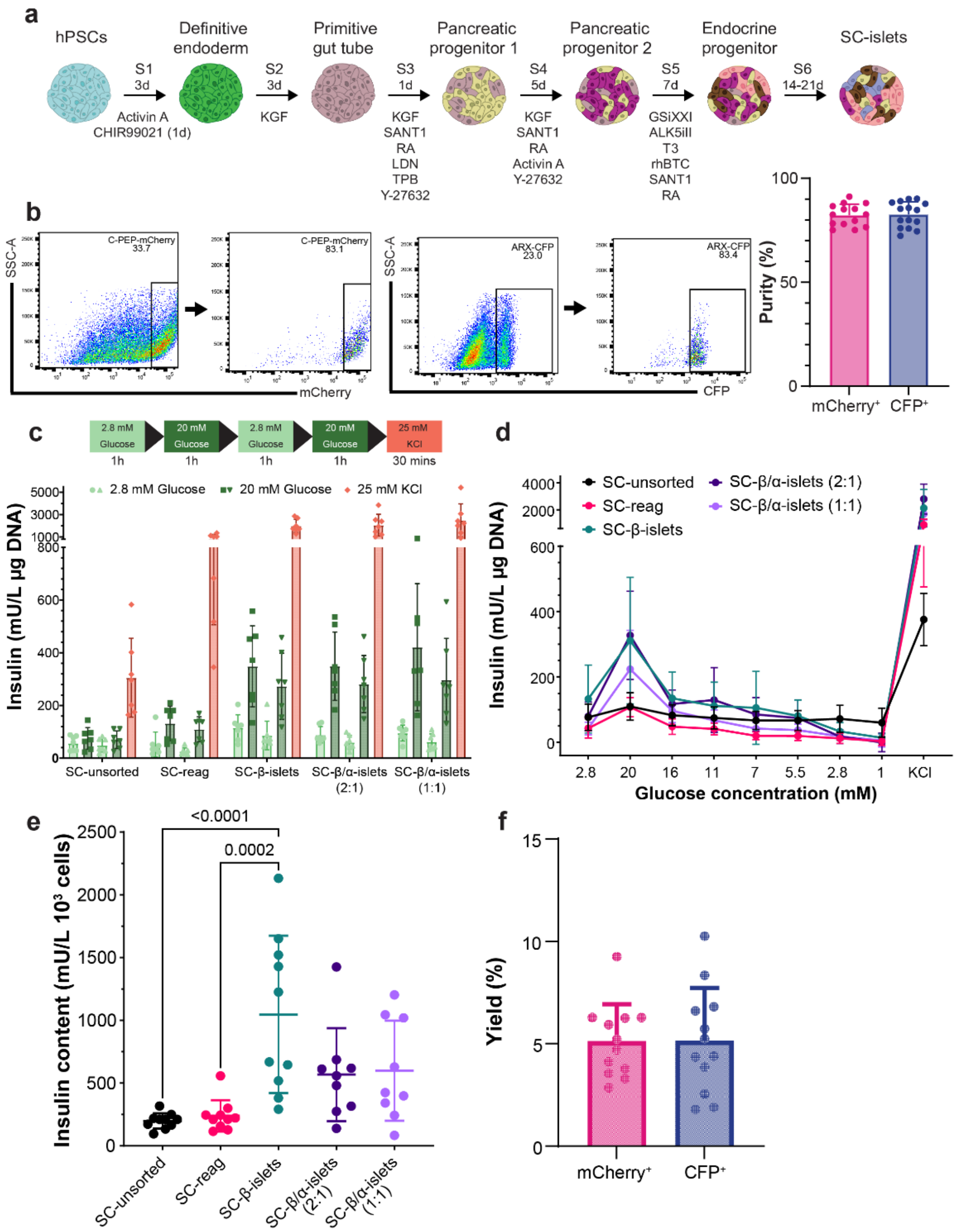


***Figure S1. ARX$^{CFP/CFP}$ x C-PEP$^{mCherry/+}$ hiPSC reporter line sorting using FACS enabled high purity sort with a low yield.*** *(a) 3D differentiation scheme protocol, consisting of 6 stages adapted from a previously published protocol[65]. (b) Representative flow cytometry analysis of sorting purity and average sorting purity of C-PEP and ARX (n=15). (c) Scheme and result of static GSIS assay normalised to DNA (n=7). (d) Insulin secretion during glucose ramp assay (n=5). (e) Amount of insulin in differently composed SC-islets normalised to $10^3$ cells. One-way ANOVA statistical test with Tukey's multiple comparison was used. (f) Amount of C-PEP-mCherry$^+$ and ARX-CFP$^+$ cells enriched using FACS in comparison to the total amount of positive cells (n=12).*

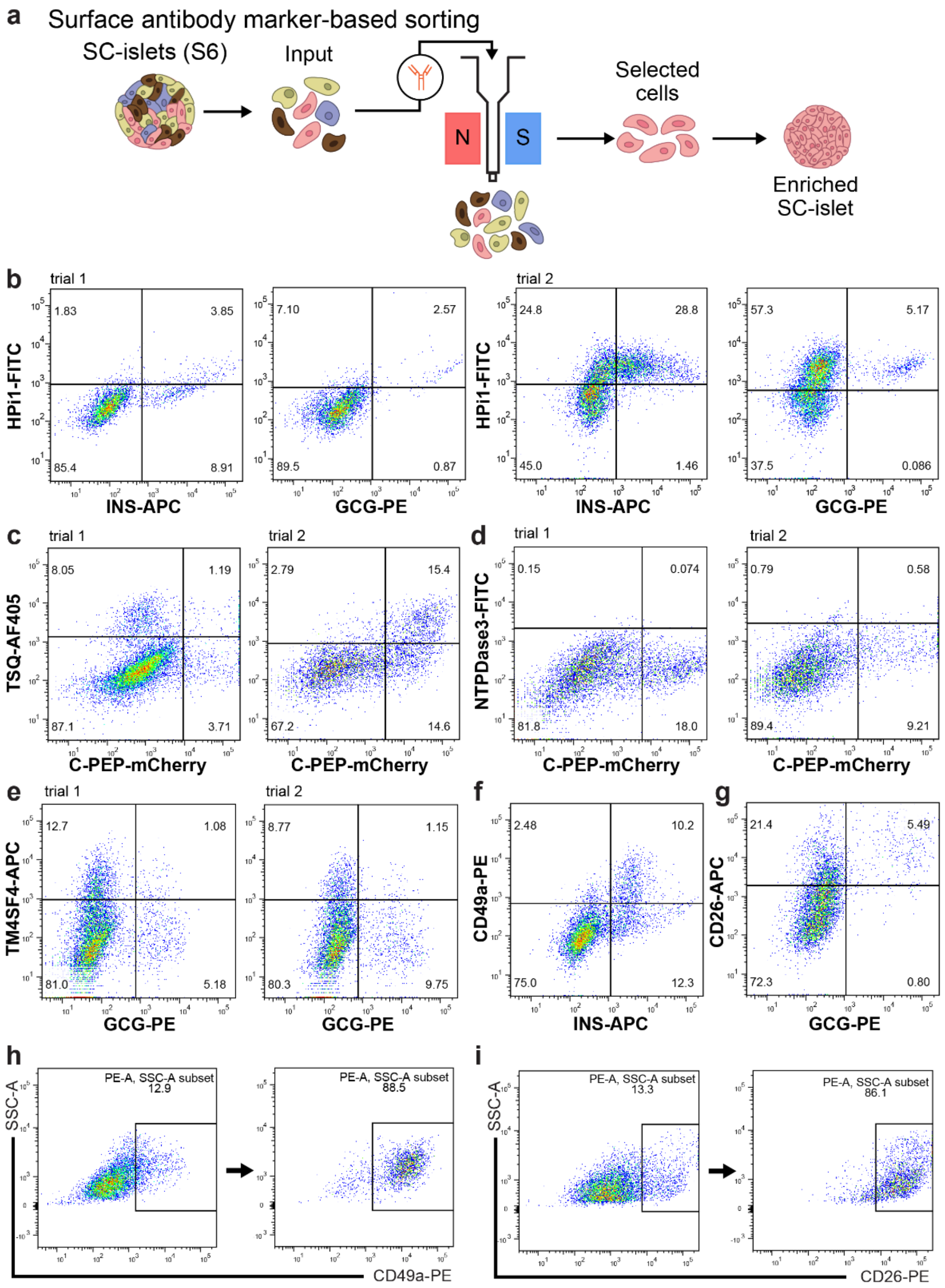


***Figure S2. Surface antibodies tested for enrichment of SC-$\beta$ and $\alpha$ -cells.*** *(a) Experimental scheme for positive selection of SC-$\beta$ and SC-$\alpha$. Flow cytometry representative plots of surface markers against SC-$\beta$- (INS or C-PEP) or SC-$\alpha$-cells (GCG). All trials were performed at various days of early-mid S6 stage (day 3-14). Tested markers are (b) pan-islet marker HPi1, (c) zinc marker TSQ, (d) $\beta$-cell marker NTPDase3, (e) $\alpha$-cell marker TM4SF4, (f) SC-$\beta$-cell marker CD49a, and (g) SC-$\alpha$-cell marker CD26. Representative flow cytometry of sorting purity using (h) CD49a and (i) CD26.*

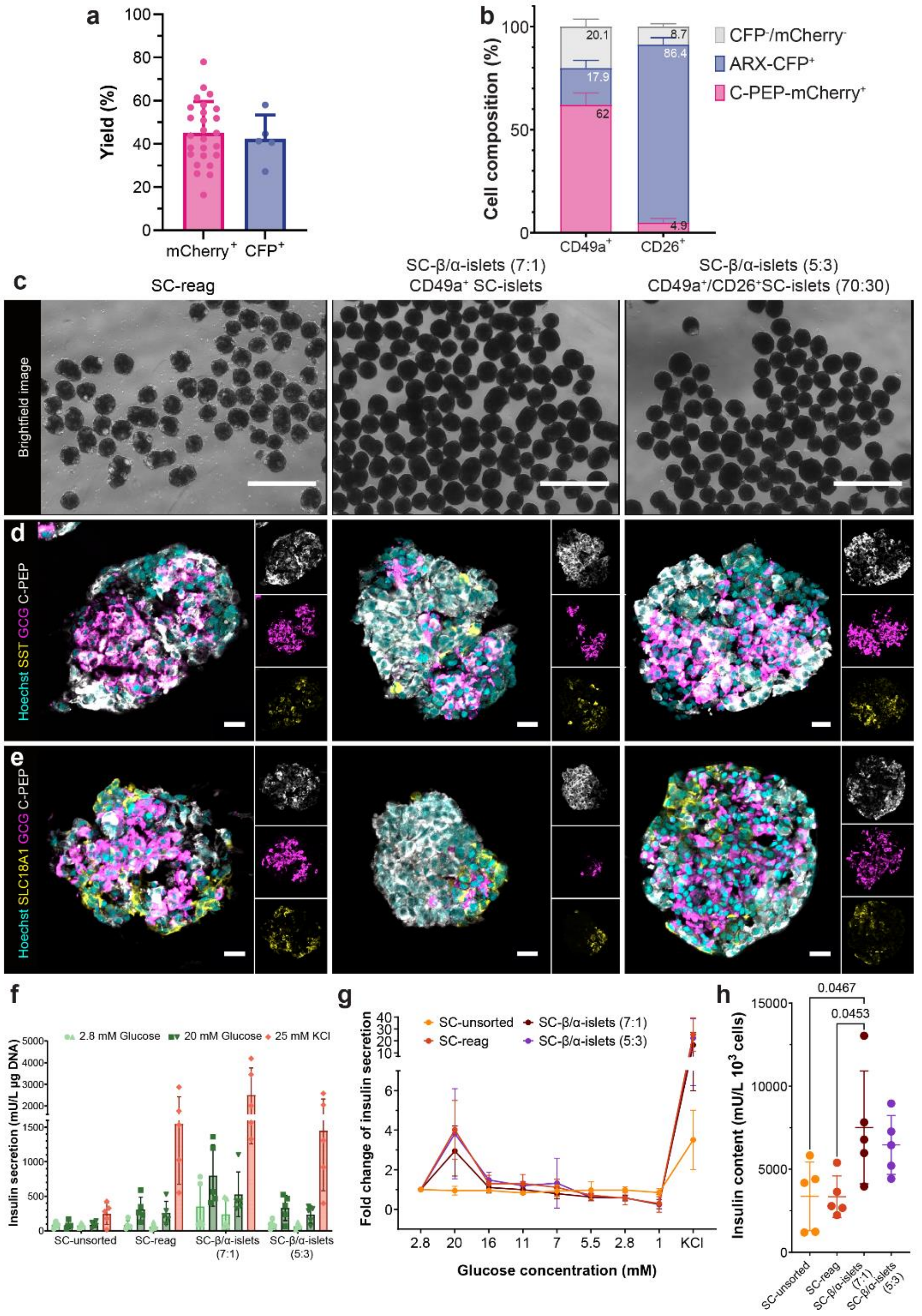


***Figure S3. Optimised sorting with CD49a and CD26 resulted in SC-$\beta$ and $\alpha$-cell enriched SC-islets with improved functionality and insulin content compared to non-enriched SC-islets.*** *(a) Amount of C-PEP (mCherry) and ARX (CFP) positive cells (measured through live fluorescence expression) enriched using MACS in comparison to the total amount of positive cells (n=5-25). (b) Cell composition of CD49a and CD26 optimised enrichment directly after sorting. (c) Brightfield image of SC-reag and enriched SC-islets 1 week after sorting. Scale bar, 750 µm. Immunostaining of (d) C-PEP, GCG, SST and (e) C-PEP, GCG, SLC18A1 encompassing all major endocrine cells found in SC-islets. Scale bar, 20 µm. (f) Insulin secretion of SC-islets upon low (2.8 mM) and high (20 mM) glucose challenge. (g) Glucose ramp result of MACS-sorted and nonsorted SC-islets. (h) Insulin content of SC-islets measured by ELISA and normalised to $10^3$ cells (n=5, One-way ANOVA statistical test* with *Tukey's multiple comparison).*

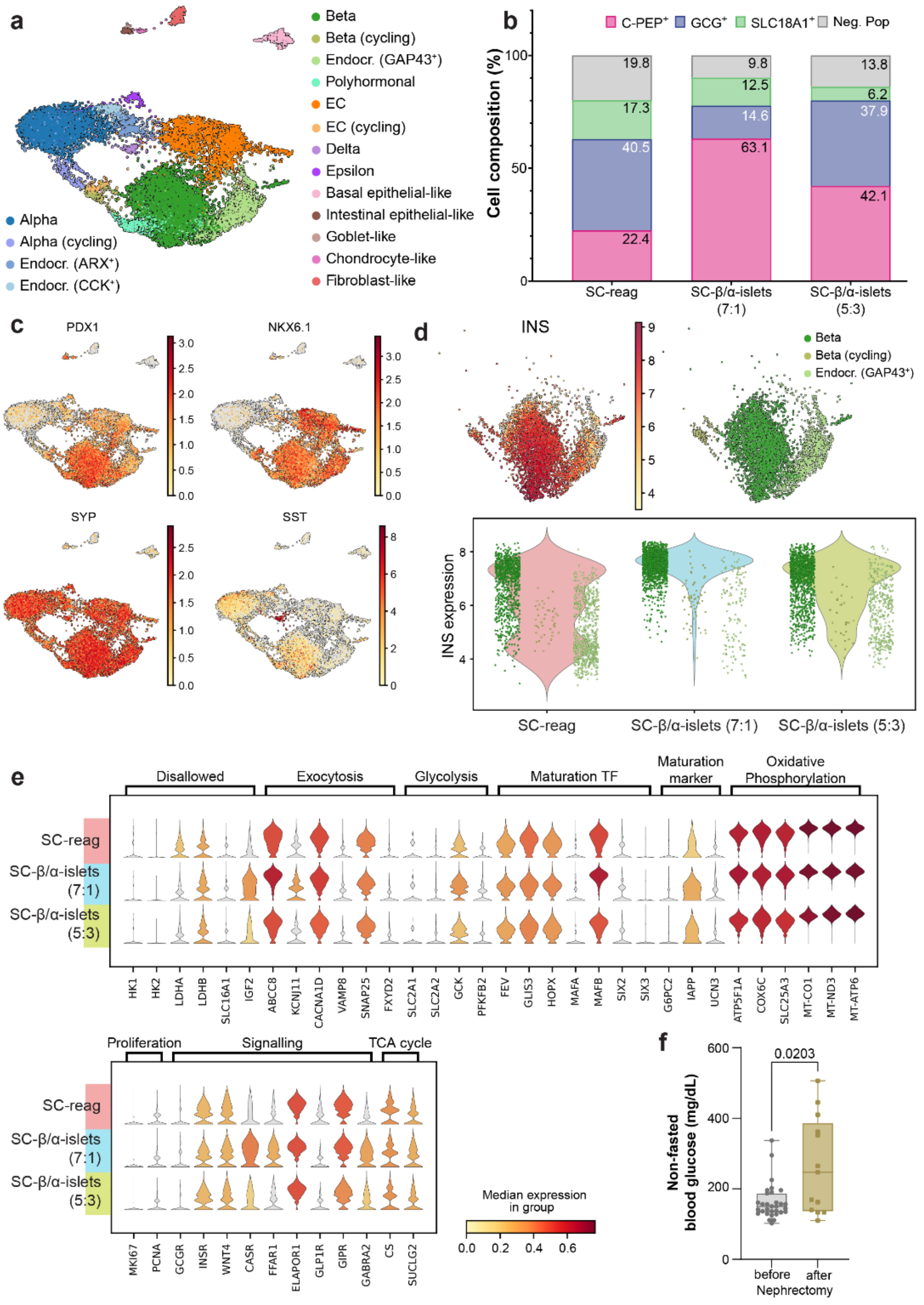


***Figure S4. CD49a enrichment increases the amount high insulin expressing cells in SC-islets.*** *(a) UMAP-based embedding coloured by fine annotation of all clusters before transplantation. (b) Cell composition based on flow cytometry analysis of scRNA-seq samples. (c) Log-normalised expression of pancreatic marker genes shown in a UMAP. (d) INS expression in beta, beta (cycling), and GAP43⁺ clusters showing the distribution of these cells in the different enrichment strategies. Bottom plot: Violin plot of imputed INS expression overlayed by scatter points coloured by cell type. (e) Stacked violin plot representation of the scaled log-normalised expression of genes associated with the function and maturation of SC-β cells by enrichment. (f) Random fed blood glucose before and after nephrectomy (min to max shown, Welch Test).*

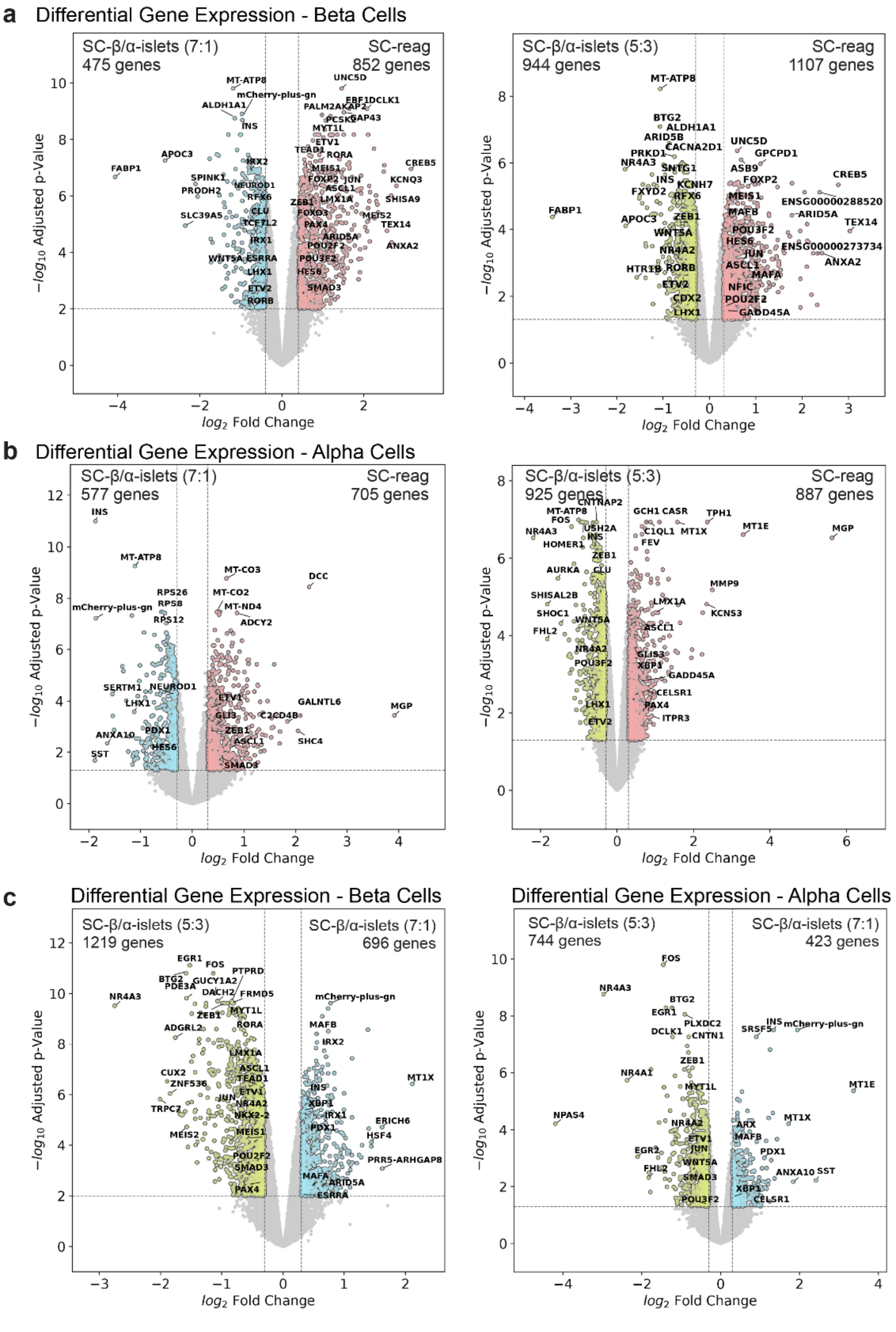


***Figure S5. Differential gene expression of SC-$\beta$ and $\alpha$-cells between different enrichment strategies before transplantation showed minimal changes.*** *Volcano plot depicting differential gene expression between enriched SC-islets vs SC-reag in (a) $\beta$-cells and (b) $\alpha$-cells, and (c) between the two enriched SC-islets in SC-$\beta$- and $\alpha$-cells.*

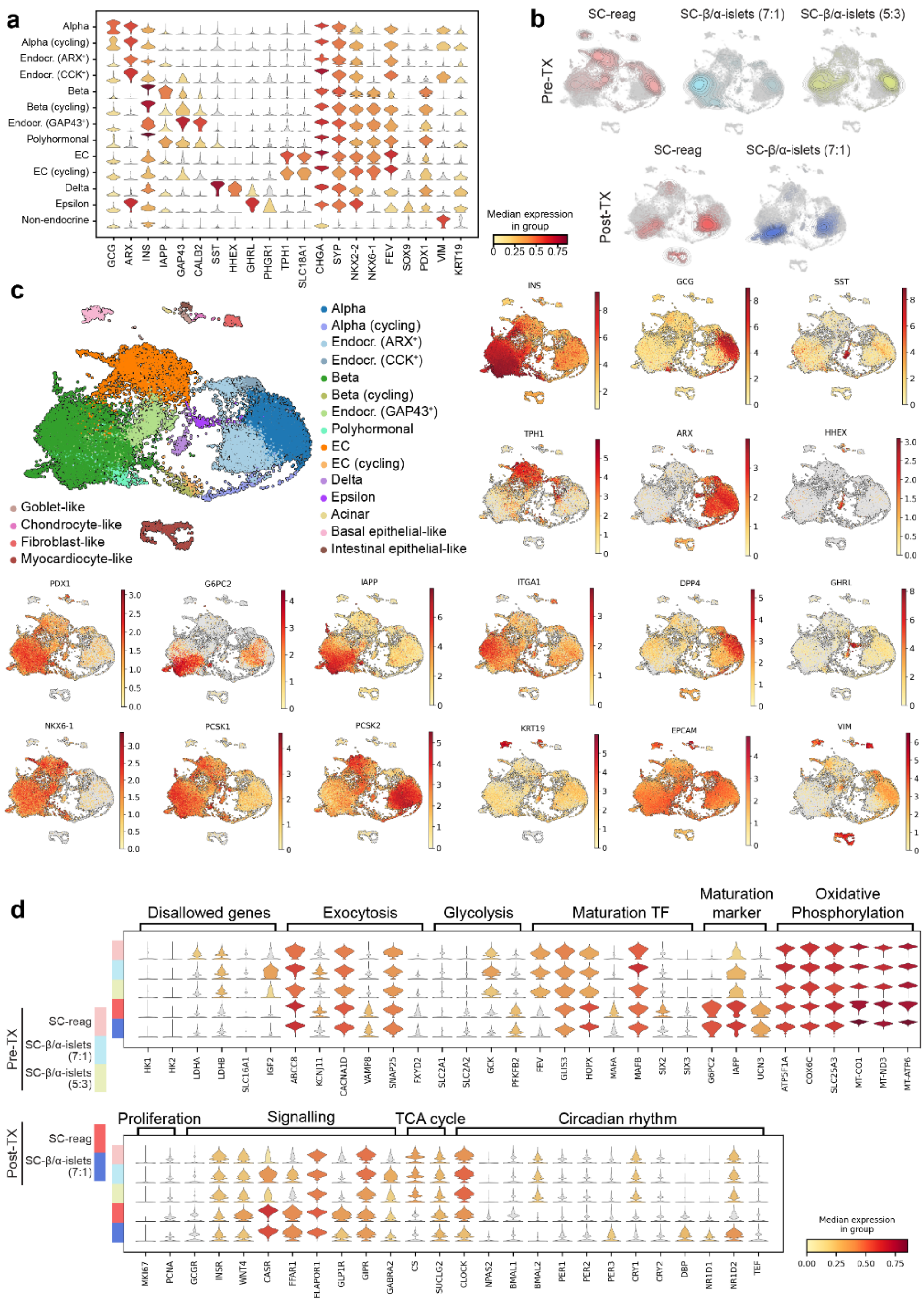


***Figure S6. Transcriptional identity of SC-β-cells matured during in vivo transplantation.*** *(a) Stacked violin plot showing the log-normalised expression of the main marker genes used for annotation. (b) Kernel density estimate plots showing the distribution of samples within the integrated UMAP. (c) Integrated UMAP with the fine annotation of all clusters before and after transplantation and log-normalised expression of gene markers for endocrine cells, pancreatic markers, β-cell maturation, key processing enzymes, ductal and epithelial/mesenchymal markers. (d) Stacked violin plot representation of genes (scaled log-normalised) associated with function and maturation of SC-β cells by each enrichment strategy and stage.*

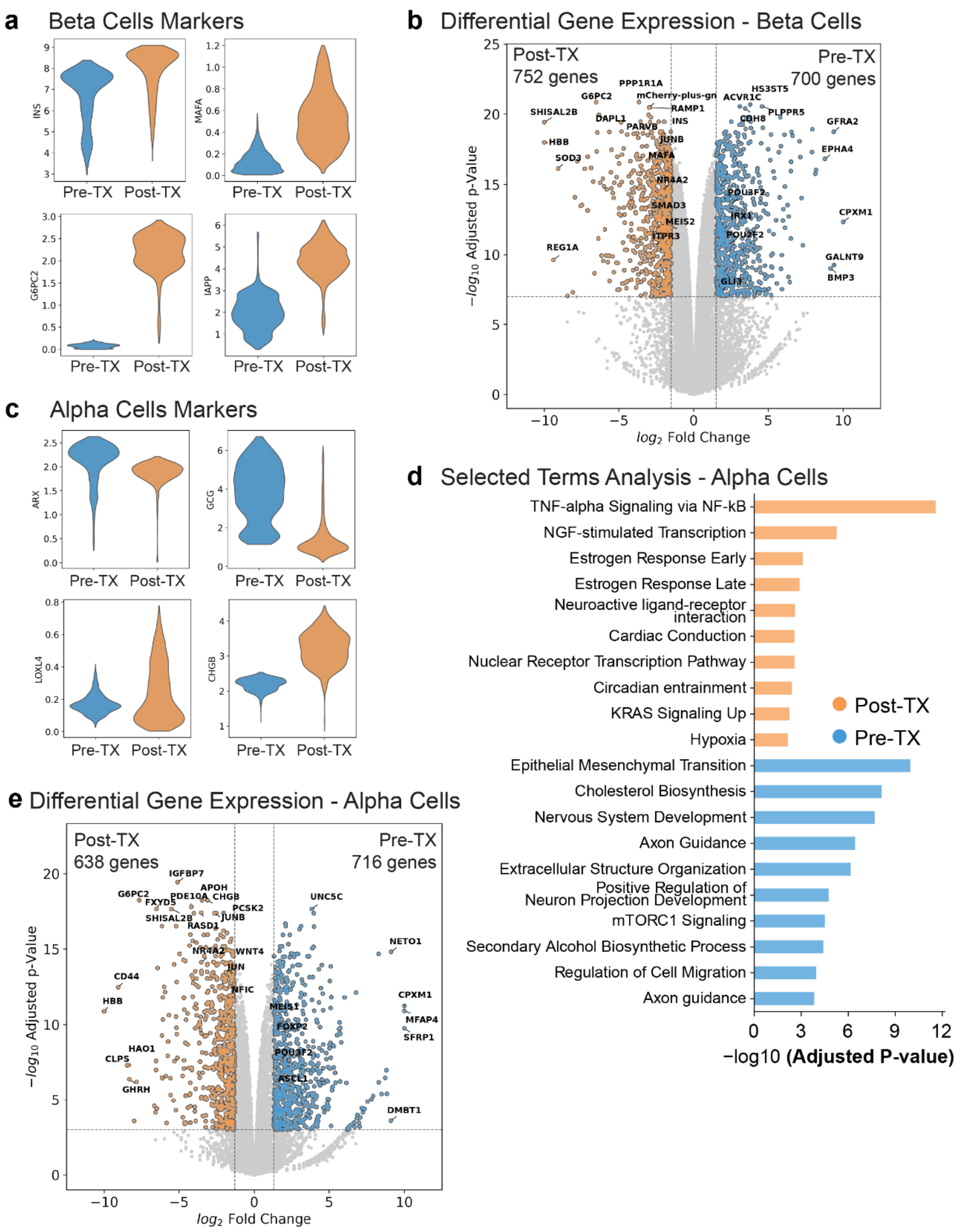


***Figure S7. SC-$\beta$ and $\alpha$-cells post-TX showed a more mature gene signature.*** *(a) Comparative violin plots of imputed expression of known $\beta$-cell markers between Pre- and Post-TX. (b) Differential gene expression of $\beta$-cells pre- vs post-TX depicted in a volcano plot. (c) Comparative violin plots of imputed expression of known $\alpha$-cell markers between pre- and post-TX. (d) Barplot of the selected pathways from the Over Representation Analysis of differentially expressed genes and (e) volcano plot of differential gene expression of SC-$\alpha$-cells before and after transplantation.*

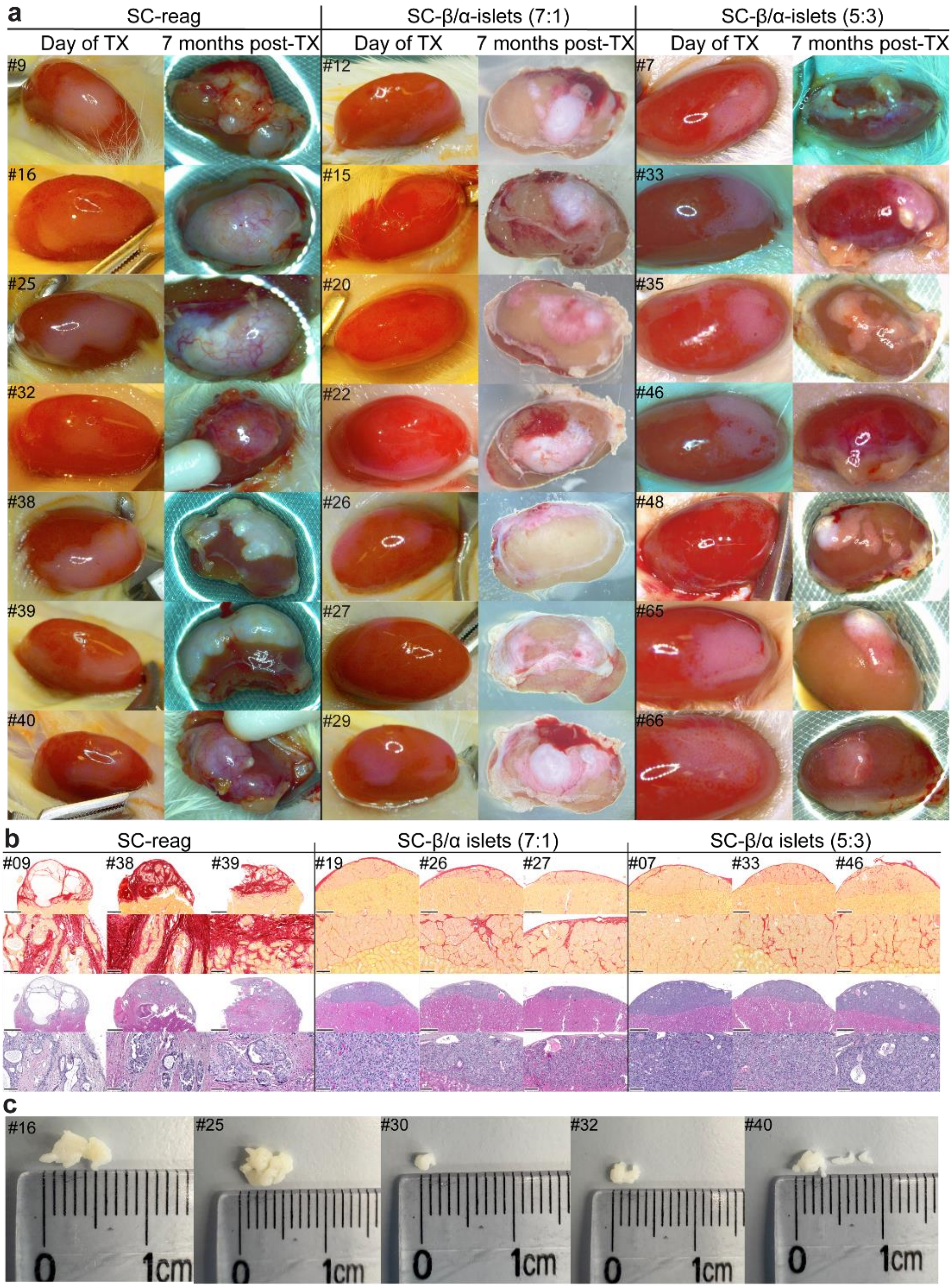


***Figure S8. Enriched SC-islets generated less outgrowth and fibrosis compared to SC-reag.*** *(a) Representative comparative image of graft on the day of transplantation (Day of TX) and the day of nephrectomy (7 months post-TX). (b) Pathological analysis of paraffin sections stained with Sirius Red (upper rows) and hematoxylin and eosin (H&E, bottom rows) staining. Scale bar, 500 µm* and *100 µm. (c) Representative images of dissected bone-like structure observed on SC-reag grafts 7 months post-TX.*

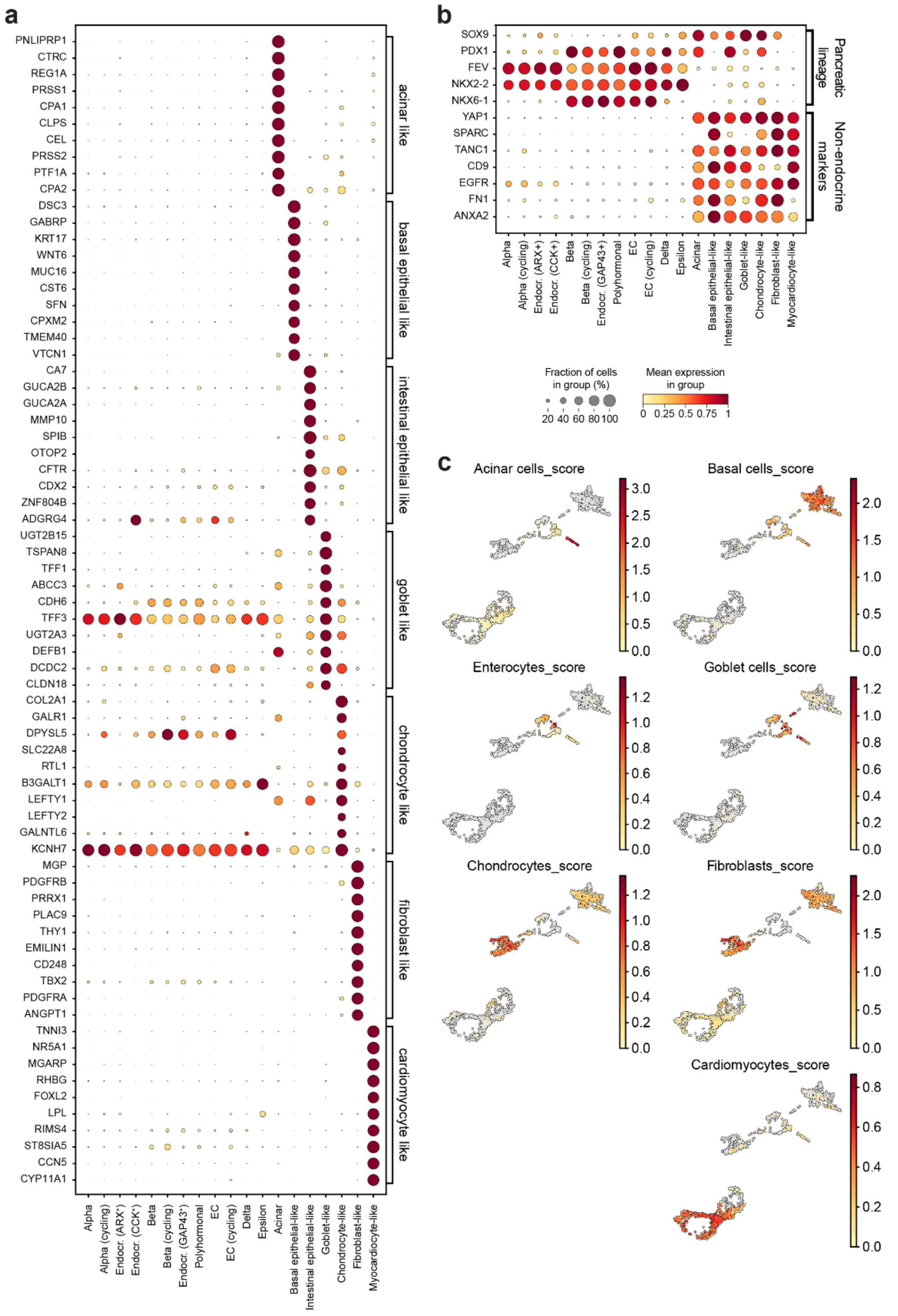

***Figure S9. Genes expressed in each of non-endocrine cell subcluster.*** *Dot plot showing the scaled log-normalised expression of (a) the top 10 marker genes for each off-target cluster and (b) comparison against the pancreatic lineage and non-endocrine markers identified. (c) UMAP-based embedding of the off-target clusters showing scores of described cell types based on the expression of associated markers found in the literature.*

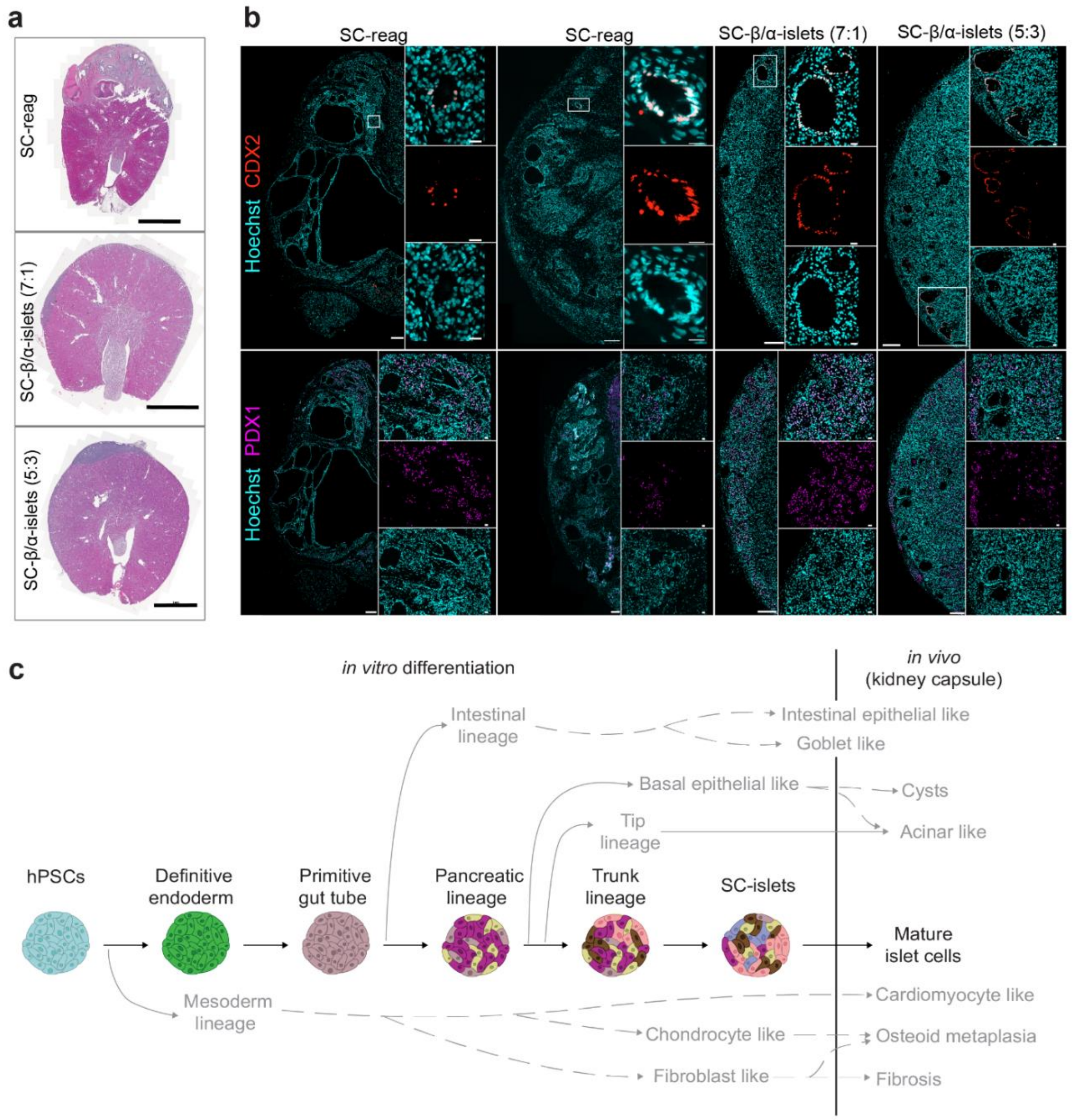


***Figure S10. Further morphological characterization of SC-islet grafts and proposed model of off-target cell origin during the differentiation.*** *(a) Whole section scan of the graft with the kidney stained with H&E staining. Scale bar, 2 mm. (b) IHC of CDX2 and PDX1 on the graft. Scale bar, 200 μm and 20 μm. (c) Proposed model of off-target source in vitro.*

*Supplementary Table 1. Differentiation protocol*

| Stage | Base media composition | Duration (day) | Compound (concentration) | Catalog No. |
|---|---|---|---|---|
| S1 | 500mL MCDB131, 0.22 g glucose, 1.23 g sodium bicarbonate, 10 g BSA, 5 mL GlutaMAX, 10 µL ITS-X, 5 mL P/S | 1 | Activin A (100 ng/mL) | Peprotech, 120-14-500 |
| | | | CHIR99021 (3 µM) | Reprocell, 25704-0004 |
| | | | Vitamin C (0.25 mM) | Merck, Sigma-Aldrich, A4544 |
| | | 2-3 | Activin A (100 ng/mL) | Peprotech, 120-14-500 |
| | | | Vitamin C (0.25 mM) | Merck, Sigma-Aldrich, A4544 |
| S2 | 500mL MCDB131, 0.22 g glucose, 0.615 g sodium bicarbonate, 10 g BSA, 5 mL GlutaMAX, 10 µL ITS-X, 5 mL P/S | 1-3 | FGF-7 (50ng/mL) | Peprotech, 100-19-100 |
| | | | Vitamin C (0.25 mM) | Merck, Sigma-Aldrich, A4544 |
| S3 | 500mL MCDB131, 0.22 g glucose, 0.615 g sodium bicarbonate, 10 g BSA, 5 mL GlutaMAX, 2.5 mL ITS-X, 5 mL P/S | 1 | FGF-7 (25ng/mL) | Peprotech, 100-19-100 |
| | | | SANT-1 (0.25 mM) | Sigma Aldrich, S4572 |
| | | | Retinoic Acid (2 µM) | Sigma Aldrich, R2625 |
| | | | LDN-193189 (200 nM) | Reprocell, (25704-0074) |
| | | | TPB (α-Amyloid precursor protein modulator; 0.5 µM) | Merck Millipore, 565740 |
| | | | Y-27632, Rocki (10 µM) | Tocris, 1254/10 |
| | | | Vitamin C (0.25 mM) | Merck, Sigma-Aldrich, A4544 |
| S4 | 500mL MCDB131, 0.22 g glucose, 0.615 g sodium bicarbonate, 10 g BSA, 5 mL GlutaMAX, 2.5 mL ITS-X, 5 mL P/S | 1-5 | FGF-7 (25ng/mL) | Peprotech, 100-19-100 |
| | | | SANT-1 (0.25 mM) | Sigma Aldrich, S4572 |
| | | | Retinoic Acid (0.1 µM) | Sigma Aldrich, R2625 |
| | | | Activin A (5 ng/mL) | Peprotech, 120-14-500 |
| | | | Y-27632 (Rocki; 10 µM) | Tocris, 1254/10 |
| | | | Vitamin C (0.25 mM) | Merck, Sigma-Aldrich, A4544 |
| S5 | 500mL MCDB131, 1.8 g glucose, 0.877 g sodium bicarbonate, 10 g BSA, 5 mL GlutaMAX, 2.5 mL ITS-X, 5 mL P/S, 5 mg heparin | 1-7 | gamma Secretase inhibitor XXI (1 µM) | Merck, 565790 |
| | | | ALK5InhibitorII (10 µM) | EnzoLifeSciences, ALX-270-445-M005 |
| | | | T3 (3,3',5-triiodo-L-thyronine sodium; 1 µM) | Sigma Aldrich, T6397 |
| | | | SANT-1 (0.25 mM) | Sigma Aldrich, S4572 |
| | | | Retinoic Acid (0.1 µM) | Sigma Aldrich, R2625 |
| | | | Betacellulin (20 ng/mL) | Bio-Techne, 261-CE-250/CF |
| | | | Vitamin C (0.25 mM) | Merck, Sigma-Aldrich, A4544 |
| S6 | 500mL MCDB131, 0.23 g glucose, 10.5 g BSA, 5.2 mL GlutaMAX, 5.2 mL P/S, 5.2 mL MEM nonessential AA, 5 mg heparin, 84 µg ZnSO4, 523 µL Trace Elements A, 523 µL Trace Elements B | 1-21 | | |

*Supplementary Table 2. Base media details*

| Base media material | Catalog No. |
|---|---|
| MCDB 131 Medium | Thermo Fisher Scientific, Gibco, 10372-019 |
| D-(+)-Glucose | Merck, Sigma-Aldrich, G7528 |
| Sodium bicarbonate ($NaHCO_3$) | Carl Roth, 8551.1 |
| BSA Fraction V, Fatty Acid Free, for cell culture media | Roche, 10775835001 |
| GlutaMAX Supplement | Thermo Fisher Scientific, Gibco, 35050038 |
| ITS-X (Insulin-Transferrin-Selenium-Ethanolamine ;100X) | Thermo Fisher Scientific, Gibco, 51500-056 |
| P/S (Penicillin-Streptomycin; 10000 U/mL) | Thermo Fisher Scientific, Gibco, 15140122 |
| Heparin | Merck, Sigma-Aldrich, H3149-10KU |
| MEM nonessential Amino Acid Solution | Corning, 15333581 |
| Zinc Sulfate Heptahydrate ($ZnSO_4 \cdot 7H_2O$) | Merck, Sigma-Aldrich, Z0251 |
| Trace Elements A, 1000x solution | Corning, 15333641 |
| Trace Elements B, 1000x solution | Corning, 15343641 |

*Supplementary Table 3. Antibody list*

| Epitope | Host | Conjugate | Dilution | Supplier | Catalog No. | Assay |
|---|---|---|---|---|---|---|
| CD26/DPP-4 | goat | - | 1:50 | R&D Systems | AF1180 | IHC |
| CDX2 [SFI-2] | mouse | - | 1:50 | DCS | CI841C01 | IHC |
| C-Peptide | guinea pig | - | 1:300 | Abcam | ab30477 | IHC |
| Cytokeratin 19 [EP1580Y] | rabbit | - | 1:200 | Abcam | ab52625 | IHC |
| Glucagon | guinea pig | - | 1:1500 | TAKARA | M182 | IHC |
| Glucagon | mouse | - | 1:100 | Santa Cruz | sc-514592 | IHC |
| Insulin | mouse | - | 1:800 | Cell signaling | 8138 | IHC |
| Integrin alpha 1 / CD49a | Sheep | - | 1:200 | R&D Systems, Inc. | AF5676-SP | IHC |
| PDGFRB | rabbit | - | 1:100 | Cell signaling | 3169 | IHC |
| PDX1 | goat | - | 1:100 | R&D Systems | AF2419 | IHC |
| RFP (5F8) | rat | - | 1:800 | chromotek | ORD003515 | IHC |
| SLC18A1 | rabbit | - | 1:200 | Atlas Antibodies | ATA-HPA063797-100 | IHC |
| Somatostatin | rat | - | 1:300 | Invitrogen | MA5-16987 | IHC |
| Somatostatin | mouse | - | 1:500 | Santa Cruz | sc-55565 | IHC |
| Synaptophysin | mouse | - | 1:200 | DaKo | M0776 | FC |
| Insulin | Mouse IgG1, k | Alexa Fluor® 647 | 1:40 | BD | 565689 | FC |
| Glucagon | Mouse IgG1, κ | FITC | 1:180 | Novus Biologicals | NBP2-21803F | FC |
| CD26 | Mouse IgG2, κ | PE | 0.05 µl/$10^6$ cells | Miltenyi Biotec | 130-126-362 | MACS/ FACS |
| CD49a, Clone SR84 | Mouse IgG1, k | PE | 0.4 µl/$10^6$ cells | BD | 559596 | MACS/FACS |
| PE-beads | | magnetic | 1 µl/$10^6$ cells | Miltenyi Biotec | 130-048-801 | MACS |

IHC= Immunohistochemistry, FC= Flow cytometry
